# Cryptic genetic variations of alanine:glyoxylate aminotransferase shape its fitness and dynamics

**DOI:** 10.1101/2021.05.24.445519

**Authors:** Mirco Dindo, Stefano Pascarelli, Davide Chiasserini, Silvia Grottelli, Claudio Costantini, Gen-Ichiro Uechi, Giorgio Giardina, Paola Laurino, Barbara Cellini

**Affiliations:** Protein Engineering and Evolution Unit, Okinawa Institute of Science and Technology Graduate University, 1919-1 Tancha, Onna, Okinawa, Japan 904-0495; Department of Medicine and Surgery, University of Perugia, P.le L. Severi 1, 06132 Perugia, Italy; Department of Biochemical Sciences “A. Rossi Fanelli”, Sapienza University of Rome, Italy

## Abstract

Genetic variations expand the conformational landscape of proteins and may underlie cryptic properties that promote environmental adaptability. However, they can also represent modifying factors for disease susceptibility, by changing frustrated regions that in turn affect protein overall intracellular fitness. In this dichotomy between conservation and innovation, understanding at structural level how genetic variations keep the balance to maintain protein fitness represents an unmet need.

Herein, we took advantage of known genetic variations of human alanine:glyoxylate aminotransferase (AGT1), which is present as a common major allelic form (AGT-Ma) and a minor polymorphic form (AGT-Mi) expressed in 20% of Caucasian population. By crystallographic studies and molecular dynamics simulations we showed that the polymorphic amino acid substitutions shape the conformational flexibility of AGT1 so that three surface regions that are structured in AGT-Ma become disordered in AGT-Mi, thanks to plasticity effects propagated from the mutation site(s) to the whole structure. In-depth biochemical characterisation of variants from a library encompassing the three regions correlate this plasticity to a fitness window between AGT-Ma and AGT-Mi, and suggest the existence of cryptic functions related to protein-protein interactions. These results establish that naturally-occurring genetic variations tip the balance between stability and frustration to expand the potential innovability of the protein.

## INTRODUCTION

Proteins might exhibit underlying functions that are not phenotypically observed under normal conditions, but they represent a genetic potential that might emerge as favourable under rapid selection conditions, a phenomenon called “cryptic” genetic variation (1). Cryptic genetic variations contribute to protein evolvability, innovability, and adaptability to cellular stresses. However, they can also influence the emergence of complex diseases as well as represent modifier factors in the phenotypic expression of rare mutations. In this regard, cryptic alleles represent a trade-off between an advantageous adaptation when an unusual condition becomes common, and the generation of polymorphic traits that increase disease susceptibility (2, 3). Is it possible to unravel the structural basis of this trade-off?

Cryptic genetic variations can have subtle effects on protein structure that are difficult to identify. Often they widen the protein conformational sampling, by increasing the degree of disorder (4). An extreme consequence of a wide conformational sampling is seen in protein regions that are conformationally frustrated and thus explore an excess of flexibility at the edge of disorder (fuzzyness), or simply don’t have a defined structure, i.e. intrinsically disordered regions (IDRs) (5, 6). This structural plasticity can be beneficial for a protein since it can mediate the interaction with multiple binding partners (7, 8). At the same time, increasing frustration and disorder negatively affects protein stability, which in turn can decrease protein fitness in a specific intracellular environment - herein we refer to protein fitness as the ensemble of properties contributing to the function(s) of a protein, such as protein’s stability and activity. However, how evolution balances cryptic genetic variations hidden behind the conformational plasticity of frustrated regions or IDRs and how they can have a repercussion on protein fitness remains poorly understood.

Liver peroxisomal alanine:glyoxylate aminotransferase 1 (AGT1) is a dimeric pyridoxal-5′ phosphate (PLP)-dependent enzyme that transaminates L-alanine and glyoxylate to pyruvate and glycine, respectively, a reaction crucial for glyoxylate detoxification (9). Numerous naturally-occurring variants of AGT1 are known, some of which are associated with the life-threatening condition named Primary Hyperoxaluria Type I (PH1) characterized by the progressive accumulation of calcium oxalate stones first in the urinary tract, and then at systemic level (10, 11). Besides pathogenic forms, it is of note that 20% of the Caucasian population express a polymorphic form of AGT1 called minor allele (AGT-Mi) differing from the more common major allele (AGT-Ma) for the P11L and I340M substitutions (12). AGT-Mi is not pathogenic *per se*, but can influence the effects of other mutations in a form of genetic epistasis (13, 14). Indeed, AGT-Ma and AGT-Mi have been proposed as two limit forms for AGT1 thermodynamic and kinetic stability, placing AGT-Mi at the edge of misfolding, so that any further destabilizing mutation leads to a decrease in protein fitness resulting in aggregation and premature degradation (15, 16). The fact that AGT-Mi it is still retained in the human genome and did not go through purging selection during evolution (14) strongly suggests that it must underlay a different function. Thus, AGT1 represents an ideal model to understand how nature compromises on structure stability to ensure potential innovability. Indeed, the tight dependency of AGT1 fitness on the cellular environment makes it an ideal protein model to understand the interplay between protein fitness and cellular functional constraints.

In this work, we have performed a systematic study of AGT1 showing that AGT-Ma retains a certain degree of conformational plasticity whose limit is represented by AGT-Mi, ultimately showing the outcome of cryptic variations at the cellular level. The structural analyses of AGT-Mi allow to unravel the plasticity of AGT1 constituted by three regions whose dynamics is the results of an enhanced conformational flexibility of the N-terminal arm that propagates through all the structure due to the presence of conformational frustration. A mainly “anti-consensus” library of AGT-Ma in these three regions reveals the biochemical and biophysical aspects related to the transition from the fitness of AGT-Ma to the lowest limit of fitness represented by AGT-Mi. Finally, studies in eukaryotic cells demonstrate that an altered conformational plasticity affects intracellular stability and interaction with binding partners, suggesting that cryptic genetic variants can shape the cellular interactome.

## RESULTS

### The crystal structure of AGT-Mi unveils three unexpected disordered regions and allows to decipher AGT1 hidden plasticity

We solved the structure of AGT-Mi by X-ray crystallography at 2.2 Å resolution (**Fig. 1; Table S1**). The I340M mutation was clearly observed as a sharp positive electron density peak corresponding to the sulphur atom position (**Fig. S1A**). On the other hand, no electron density was observed for the N-terminal residues (residues 1 to 10: MASHKLLVTP) (**Fig. S1B**), indicating that the P11L mutation results in a loss of defined structure in this region. In the AGT1 dimer the N-terminal arm is involved in an important interaction between the monomers, and mutations altering its conformation are known to affect protein thermodynamic and kinetic stability (16–18). Thus, our data provide a definitive proof for many biochemical and *in silico* studies performed in the last decade, which have suggested that the lower stability of AGT-Mi was linked to a disordered N-terminus (18–22). Surprisingly, we found the presence of another disordered region in AGT-Mi: an undefined electron density was observed for residues 97-102 (GDSFLV), 121-146 (ARVHPMTKDGGHYTLQEVEEGLAQH) and 170-177 (GELCHRYK), which were not included in the final model **(Fig. S1 C, D)**. In AGT-Ma these regions form two α-helices and two β-strands at the periphery of the large domain (the structural unit binding PLP) (**Fig. 1A**) that tend to display a higher mobility in all the structures solved to date, with temperature factors (B-factors) that appear systematically higher than the mean values of each structure (**Fig. 1 B, C**). Thus, AGT1 samples different conformations whose relative populations is driven by the presence of two polymorphic changes. Ultimately, these mutations induce an increased disorder not only at the N-terminus, but also in structurally frustrated regions of AGT1. The finding that AGT-Mi is characterized by the presence of IDRs is extremely important since IDRs in globular proteins consist of the terminal regions or represent interdomain linkers (23). Usually, the shift between order and disorder is observed as a consequence of numerous factors including ligand or partner binding, while shifts caused by naturally-occurring polymorphic mutations have not been widely explored (8, 24).

**Figure 1.**
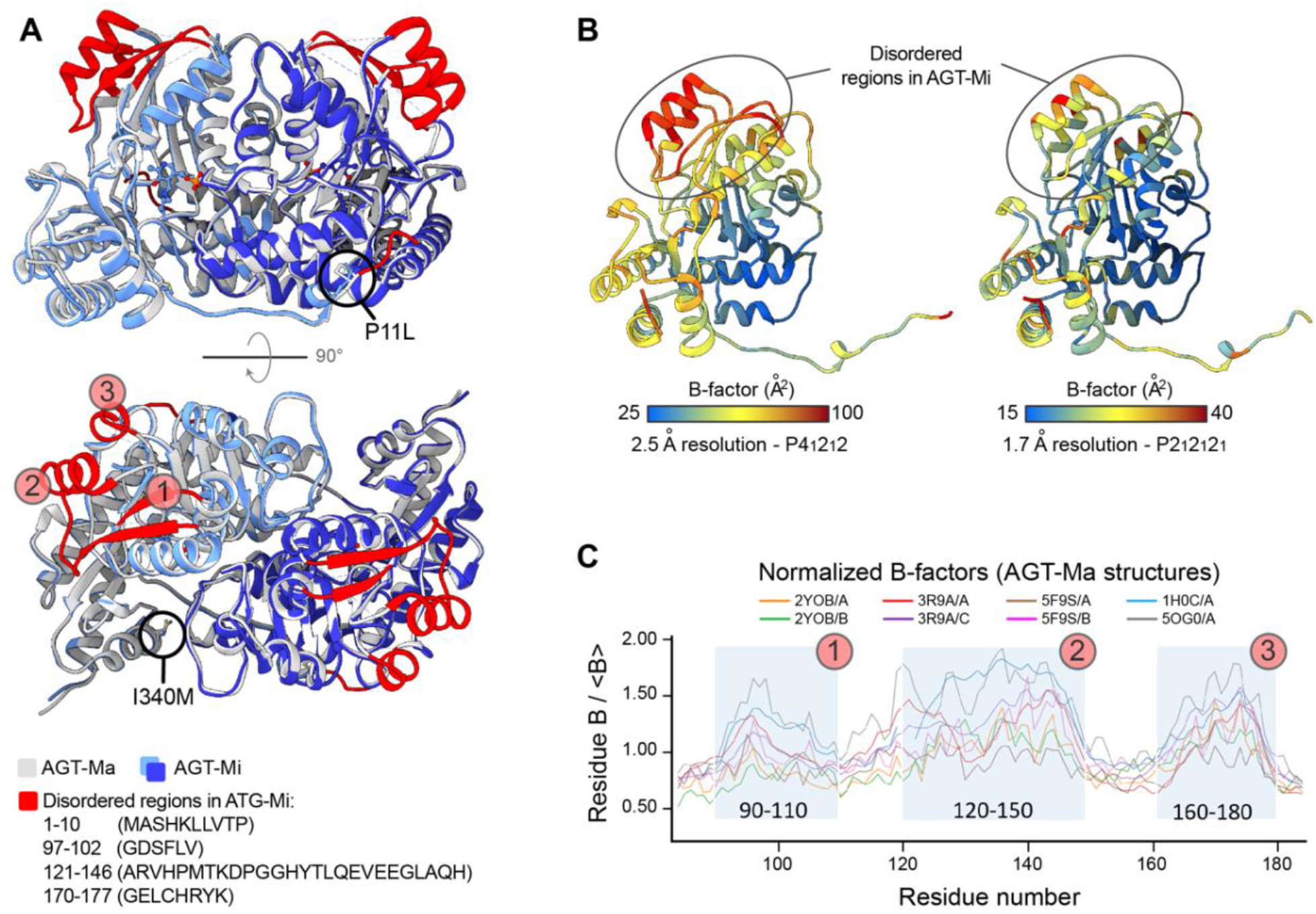
Structure of AGT-Mi. **A**) Cartoon representation of the structure of AGT-Mi superimposed on that of AGT-Ma (PDB: 5F9S). The two subunits of AGT-Mi are shown in dark and light blue while for AGT-Ma are shown in white and dark grey. The regions that were not included in the model of AGT-Mi, due to disorder, are coloured in red in the structure of AGT-Ma. Except for the N-terminus, the other disordered regions are numbered from 1 to 3 and correspond to residue stretches 97-102, 121-146 and 170-177, respectively. **B**) Two AGT-Ma structures solved at different resolution and in different space groups are coloured according to temperature factor (B-factors) values (PDBs: 5OG0 and 5F9S on the left and right side, respectively). Both structures show high B-factors in the region that are disordered in AGT-Mi. **C**) Analysis of normalized B-factors over the range of all available AGT-Ma structures in different space groups. High B-factors values correspond to the disordered regions (1, 2 and 3) in AGT-Mi. These regions are not involved in crystal contacts; therefore the observed disorder is not a consequence of crystal packing. The B-factor of the structure 2YOB is the lowest, suggesting a destabilizing effect of the P11L mutation balanced by a stabilizing effect of the I340M mutation.

### Testing the plasticity of IDRs by Molecular Dynamics allow to study the role of these regions in AGT1

Next, we investigated the AGT1 conformational sampling by molecular dynamics (MD) simulations. We aimed at detecting any difference between the conformers sampled by the PLP-bound AGT-Ma and AGT-Mi. To be unbiased we used as a model the stable AGT-Ma conformation (PDB:5F9S) as a starting point for the single- and double-mutant models. The single-mutant models were obtained by individually mutating P11L or I340M, while the double-mutant model was obtained by mutating both positions together, to achieve the AGT-Mi model (more details in the Methods). After an equilibration step, the models were run for 200 ns in five repeats (**Fig. 2A**).

**Figure 2:**
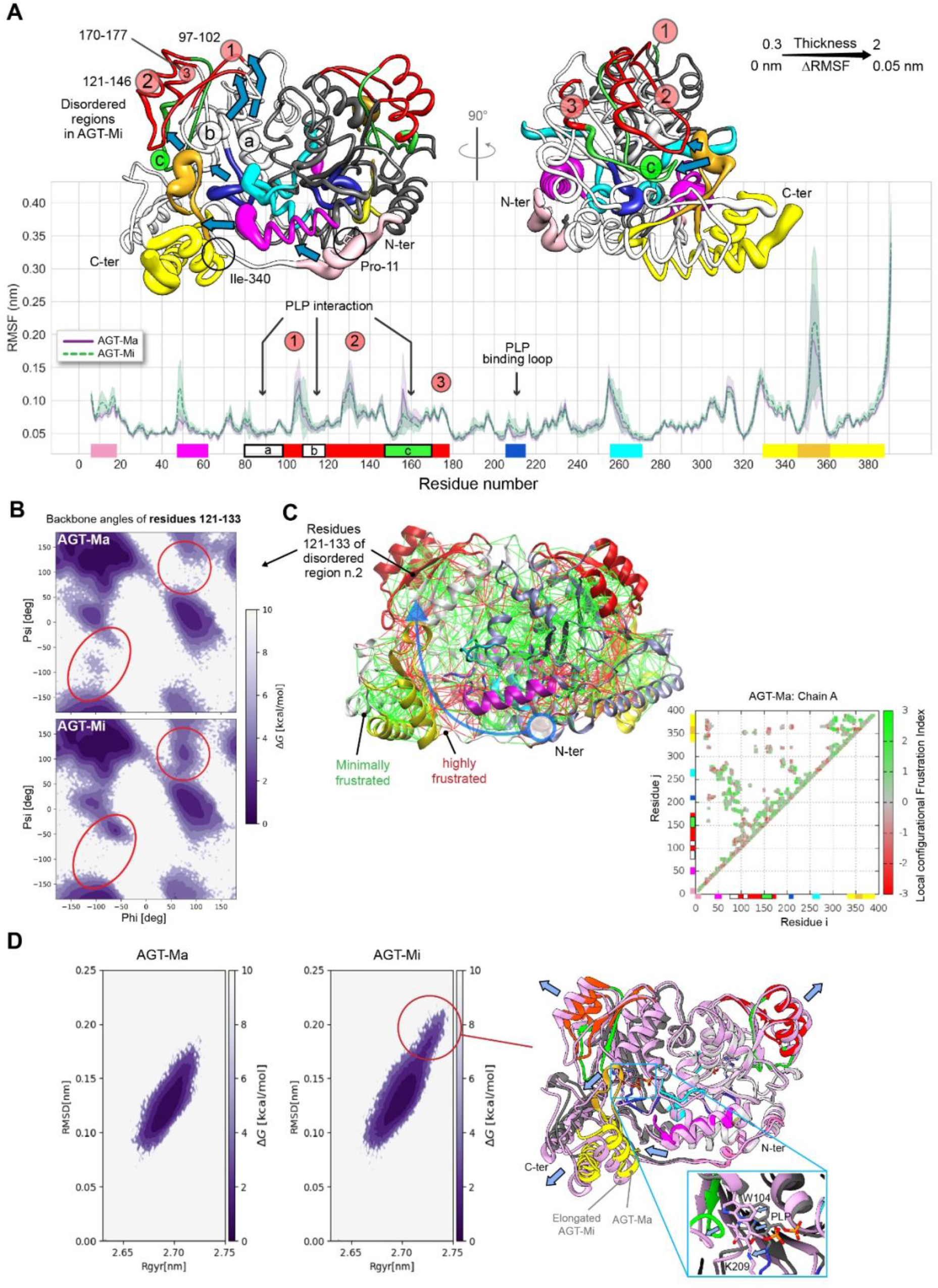
Molecular dynamics simulations. **A**) Plot of the root mean square fluctuation (RMSF) as a function of residue number: AGT-Ma (purple) and AGT-Mi (green). The difference ΔRMSF (RMSF AGT-Mi – RMSF AGT-Ma) is mapped on the structure of AGT-Ma as thickness of the tube-like presentation. The RMSF values used were obtained as the average of five trajectories. The regions of interest are marked with the same colours in the structure and in the plot. The increased mobility of the N-terminus (in *pink*) and especially of the residues 48-64 (in *magenta*) of AGT-Mi may cause a domino effect that is propagated through the C-terminus (in *yellow*) and three structural regions (indicated with *a*, *b* and *c*) that are directly interacting with PLP and with the disordered regions 1, 2 and 3 (in *red*). Details on the MD method are reported in Table S2. The structures were obtained by Chimera (26) **B**) Free energy landscape as a function of the Φ and Ψ dihedral angles of the backbone of the high B-factor region 121-133. This region forms a β-strand that contacts the flexible C-terminus. **C**) AGT-Ma conformational frustration plot. Green and red colours indicate couples of minimal and maximal frustrated residues, respectively. The configurational frustration index evaluates how the energetic contribution to protein stability of the interaction between residues *i* and *j* compares to different possible local environment. The interaction is considered frustrated when other conformations with higher stabilization energy are possible (27). Areas of high frustration are localized at the interface of the highly fluctuating regions, connecting the N-terminus to the disordered regions in ATG-Mi (blue arrow). The same colouring of panel A is used to highlight the different portions of AGT in the structure and in the correlation plot. **D**) Free Energy Surface (FES) of the five combined simulations for AGT-Ma and AGT-Mi. The first 5ns of each simulation were removed from the computations to remove any bias due to the equilibration of the system. The FES was generated in a similar fashion as in Strodel *et al.* (28) and using RMSD and gyration radius as the two coordinate systems. On the right, a representative structure of AGT-Mi in the highlighted region of panel B (with high RMSD and high gyration) is shown in pink, superposed with the structure of AGT-Ma (5F9S) that is depicted with the same colouring pattern of panel A. The loss of secondary structure and displacement (blue arrows) is evident in parts of the disordered regions and in the C-terminal domain. The displacement also affects the cofactor position (inset).

The simulations confirmed that the stretches of residues identified as IDRs in AGT-Mi by crystallographic studies also fluctuate in AGT-Ma (**Fig 2A**), as suggested by B-factor analysis (**Fig. 1B–C**), indicating that these regions are prone to disorder (frustrated) in AGT-Ma too. However, in AGT-Mi the N-terminus (residues 6-18) and the helix 48-64 quickly sample a range of alternative conformations due to an increase of the fluctuations in these regions that is not observed in the AGT-Ma simulations (**Fig. 2A**). Notably, this helix is in close contact with both the very dynamic C-terminal domain and several regions involved in PLP binding (**Fig 2A**). From the simulation results, it appears that the multiple conformations adopted by AGT-Mi result from backbone dynamics initiated by an increased fluctuation of the helix 48-64 and the N-terminus, which propagates through the C-terminus and the active site to the IDRs (**Fig. S2A**). Indeed, in the simulation we observed a difference in the distribution of the dihedral angles of backbone in second region (positions 121-131) that is the β-strand in contact with the highly flexible C-terminus (**Fig. 2B**) as well as in the region 170-177 (Fig. S2E). However, the same analysis on the other frustrated regions is not able to discriminate between AGT-Mi and AGT-Ma (Fig. S2E). Analysis of protein local frustration, nicely correlates with this picture, showing that in AGT-Ma regions of high conformational frustration cluster along the same route indicated by maximal fluctuation and connecting the N- and C-terminus dynamics (**Fig. 2C** and **movie S1**). Interestingly, Free Energy Surface (FES) analysis confirmed that AGT-Mi uniquely explores a set of conformations at high RMSD and gyration radius where the C-terminal helices and parts of the IDRs are notably displaced (**Fig 2D**). In the IDRs, this apparently small detachment of the two helices may indeed weaken the hydrophobic interactions that stabilize the structured state leading to a disordered conformation. Furthermore, MD simulations of the single mutant P11L showed enhanced fluctuations of the N-terminal tail (**Fig. S2B**), and a higher occupancy of the conformations at high radius of gyration (**Fig. S2C**). This result highlights the structural determinants of the stabilizing effect of the I340M mutation that we observed in the MD simulation of this single mutant and also in previous studies (25). In fact, Ile340 is placed at the interface between helix 48-64 and the C-terminus and its substitution with a longer hydrophobic side chain partially reduces the dynamics of these regions (**Fig S2D**).

To test the role of the order/disorder transition in a cellular environment, we rationally introduced four mutations - L101A, H146A, L172A, and C173A – that should enhance the plasticity of the IDRs by reducing the number of favourable interactions between the side chains of these regions. As shown in **Fig. S3**, substitutions in the first and third region (residues 97-102 and 170-177, respectively) do not significantly change AGT1 specific activity and protein level, while the H146A mutation of the second region leads to a significant reduction of soluble protein and active enzyme levels, possibly related to the increased backbone flexibility of adjacent regions (121-131) suggested by the MD analysis.

Overall, by combining the structural, MD and cellular data, we highlighted the presence of a structural hotspot that appears as an order-disorder conformational switch for AGT1 in the native state. These results place AGT-Ma and AGT-Mi at the boundaries of an unexpected conformational landscape that is dominated by the equilibrium between order and disorder of the IDRs, with the major allele sampling, on average, ordered conformations and the minor allele *living* on the edge of frustration and disorder.

### A library of the disordered regions of AGT1 hints to the advantage of structural instability

To better define the role of the three most flexible regions (residues 97-102; 121-146; 170-177) on the conformational sampling of AGT1, we generated a library of AGT-Ma variants in these regions. The design of the library was based on two main principles. First, we identified all the positions where the residue conservation is below 80% among the alignment (**Table S3**). Then we identified “anticonsensus” mutations, namely those causing the change of a residue with another one present in nature in the same position but showing a lower abundance (**Fig. 3A**). This choice has been dictated by the need to mimic a subtle destabilization typical of cryptic genetic variations, while avoiding big conformational changes and keeping intact the AGT1 overall structure. Positions 127 and 177 are the only ones in which the human sequence does not show a consensus residue. For these two residues, we decided to include also mutations to the residue most represented in other species, with the aim of exploring the possibility of achieving a more stable and active structure at the expenses of the above detected structural plasticity or flexibility.

**Figure 3.**
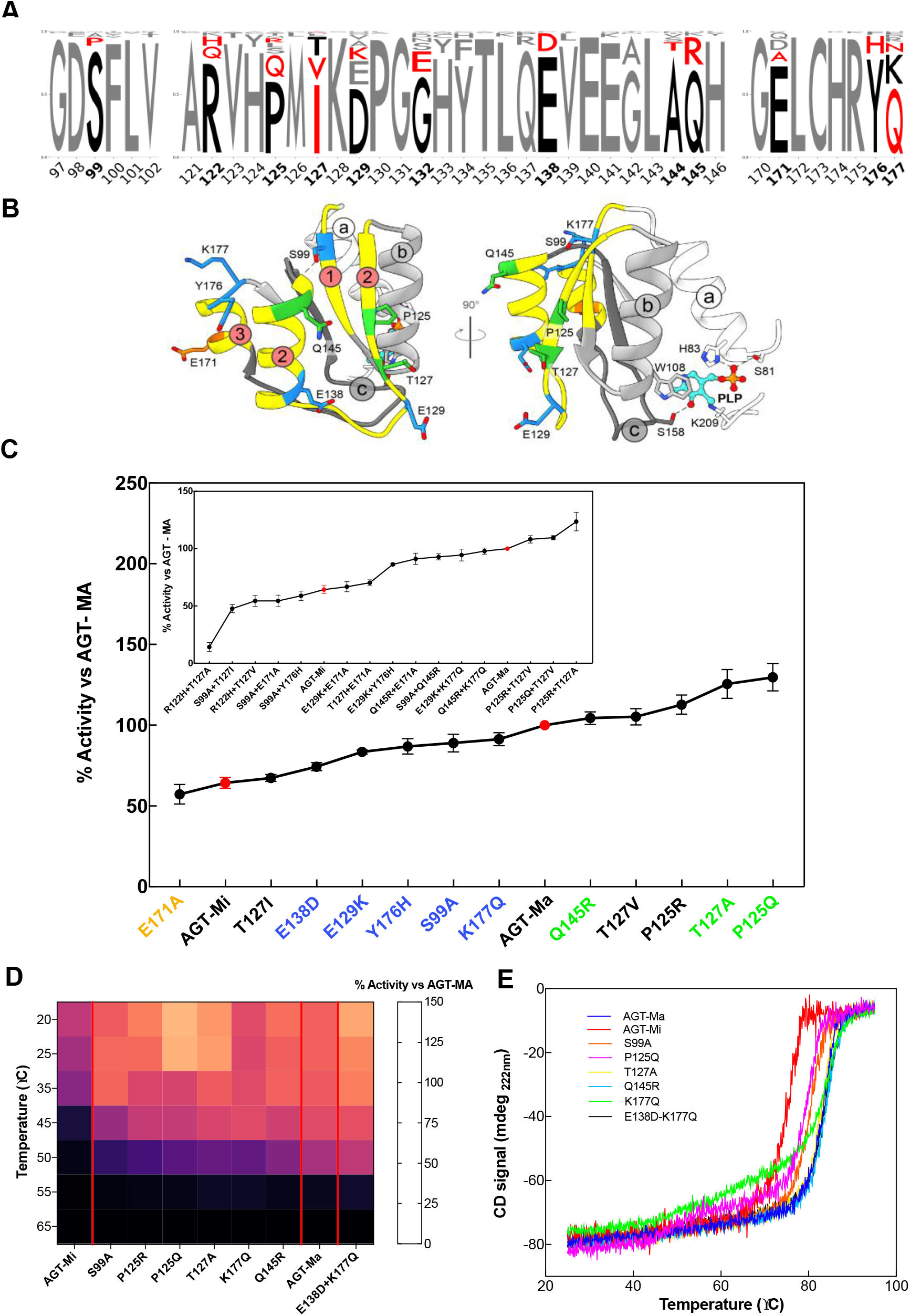
Library construction and biochemical and biophysical characterisation of the variants. **A**) Consensus logo of the three disordered regions under study. The sequences (110-130) from the non-redundant database were collected by blast search and aligned using MUSCLE algorithm on Jalview (29). The logo was generated using Logomaker (https://logomaker.readthedocs.io/en/latest/). In bold black numbers, the positions that were mutated. In black, the amino acids of the human AGT1 sequence, while in red, the aminoacids considered for the library. **B**) The mutated residues are plotted over the AGT-Ma structures in different colours. In orange the one with activity below AGT-Mi, in blue those with activity between AGT-Mi and AGT-Ma, and in green those above AGT-Ma. **C**) Residual activity of the indicated single variants as compared to AGT-Ma. Inset panel C) Residual activity of the indicated double variants as compared to AGT-Ma. Experiments have been performed in duplicate and for each single experiments three technical triplicates have been used. Data are represented as mean values ± SD. **D**) Heat map of residual activity of selected mutants obtained after incubating the lysate at different temperatures for 10 minutes. The color code is the same of panel C. **E**) Thermal stability curves of the selected variants represented in panel D. Color code is the same of panel C. Experiments have been performed in duplicate and for each single experiments three technical triplicates have been used.

Second, we used a semi-rational approach, i.e. a strategy based on the random combination of rationally-chosen mutations (see Material and Methods). As a result, we obtained a small library (48 variants) bearing single, double and triple mutations. Although the library was focused on a region distant from the catalytic site (> 12 Å, **Fig. 3B**), its importance for the overall AGT1 stability allowed the reconstruction of the putative functional intermediates within and beyond the fitness window defined by AGT-Ma and AGT-Mi (**Fig. 3C** and **Fig. S4**). 21% of the mutants showed specific activity below AGT-Mi, 48% between AGT-Mi and AGT-Ma and 31% above AGT-Ma. The wide range of activity sampled by this library demonstrates the importance of these three regions on the overall structure and gives further information on the effect of the frustration on AGT1. Nonetheless, the effects of these mutants are subtle, a sign suggestive that cryptic variations can shape the order/disorder equilibrium maintained in AGT-Ma.

To explain the fitness outcomes of the mutations introduced on the surface of the disordered regions, we chose two variants with specific activity between AGT-Mi and AGT-Ma (S99A, K177Q) and four with specific activity above the one of AGT-Ma (P125Q, T127A, Q145R, E138A/K177Q) for further characterization in the purified form. Upon confirming by CD studies that none of the introduced substitutions caused gross structural changes in AGT1 (**Fig. S5A**), we determined both the melting temperature (**Fig. 3D**) and the kinetic parameters for the transamination reaction, to get insights on possible alterations of global stability and intrinsic functional activity, respectively. The functional characterization of the purified selected variants revealed no significant changes in the kinetic parameters as reported in **Table 1** and **Fig. S6**. In fact, disordered regions are distant from the catalytic site (> 12 Å), and the mutations do not directly influence the microenvironment of the cofactor, as shown by the visible bands of the CD spectra of the corresponding variants (**Fig. S5B**). As further confirmation to this observation, we also found that the tertiary structure of the variants (**Fig. S5B**) is identical to that of AGT-Ma. More interestingly, none of the mutations interfered with the turnover number of the enzyme, thus indicating that the observed changes in specific activity in the bacterial lysate (**Fig. 3C**) can be mainly ascribed to structural alterations that affect protein levels, possibly implying that disordered surfaces represent a critical region for the overall AGT1 fitness in a cellular environment.

**Table 1.** Kinetic parameters of AGT-Ma, AGT-Mi and the variants chosen from the AGT library. Experimental details are given in the Material and Methods. Experiments have been performed in triplicate.

| Enzyme | Substrate | Cosubstrate | $k_{cat}$ ($s^{-1}$) | $K_M$ (mM) | $k_{cat}/K_M$ ($s^{-1}mM^{-1}$) |
| --- | --- | --- | --- | --- | --- |
| **AGT-Ma<sup>A</sup>** | L-alanine | Glyoxylate | $45 \pm 2$ | $31 \pm 4$ | $1.4 \pm 0.2$ |
| | Glyoxylate | L-alanine | $45 \pm 3$ | $0.23 \pm 0.05$ | $196 \pm 44$ |
| **AGT-Mi<sup>B</sup>** | L-alanine | Glyoxylate | $33 \pm 5$ | $28 \pm 2$ | $1.2 \pm 0.2$ |
| | Glyoxylate | L-alanine | $37 \pm 1$ | $0.22 \pm 0.01$ | $168 \pm 8$ |
| **S99A** | L-alanine | Glyoxylate | $36 \pm 2$ | $29 \pm 4$ | $1.2 \pm 0.18$ |
| | Glyoxylate | L-alanine | $34 \pm 3$ | $0.31 \pm 0.05$ | $109 \pm 20$ |
| **P125Q** | L-alanine | Glyoxylate | $43 \pm 5$ | $55 \pm 8$ | $0.78 \pm 0.14$ |
| | Glyoxylate | L-alanine | $35 \pm 2$ | $0.22 \pm 0.03$ | $159 \pm 22$ |
| **T127A** | L-alanine | Glyoxylate | $42 \pm 3$ | $33 \pm 4$ | $1.27 \pm 0.17$ |
| | Glyoxylate | L-alanine | $34 \pm 4$ | $0.16 \pm 0.03$ | $212 \pm 44$ |
| **Q145R** | L-alanine | Glyoxylate | $47 \pm 4$ | $41 \pm 5$ | $1.15 \pm 0.17$ |
| | Glyoxylate | L-alanine | $38 \pm 4$ | $0.21 \pm 0.04$ | $181 \pm 32$ |
| **K177Q** | L-alanine | Glyoxylate | $37 \pm 3$ | $35 \pm 5$ | $1.05 \pm 0.17$ |
| | Glyoxylate | L-alanine | $36 \pm 4$ | $0.27 \pm 0.06$ | $133 \pm 33$ |
| **E138D/K177Q** | L-alanine | Glyoxylate | $44 \pm 4$ | $53 \pm 7$ | $0.83 \pm 0.13$ |
| | Glyoxylate | L-alanine | $39 \pm 3$ | $0.27 \pm 0.04$ | $145 \pm 24$ |

<sup>A</sup>, from ref(9)
<sup>B</sup>, from ref(30)

The analysis of the melting curves of the single variants confirmed how naturally AGT1 is exploring a wide window of stability (18, 19). The stability of AGT-Ma was not affected or slightly improved by the T127A, Q145R and K177Q mutations, but it was reduced by the S99A and P125Q substitutions to values between that of the two polymorphic forms. Notably, the denaturation curves of the P125Q and K177Q variants show the loss of cooperativity on the unfolding of the two domains of AGT1, in line with MD results indicating that IDRs on the large domain influence the conformation of the small C-terminal domain where Ile340 is located (26). The latter effect on the single K177Q variant is abolished in the E138D/K177Q variant, in agreement with the improved fitness of the double variant upon *E.coli* expression (**Fig. 3C**). Overall, all the mutants included in this library are functionally active, meaning that mutations of the IDRs do not have substantial negative effects on the AGT1 structure. The finding that all the detected changes in stability and activity are subtle suggests that naturally occurring mutations can be easily tolerated while maintaining a fine fitness balance of the protein.

### Expression of IDRs variants in a cellular system highlights their role for AGT1 fitness and interactions

To investigate the effects of the library changes in a mammalian cellular environment, selected variants with single or double mutations in the disordered regions were expressed in Human Embryonic Kidney 293 (HEK293) cells and analysed for specific activity and protein levels (**Fig. 4**).

**Fig. 4:**
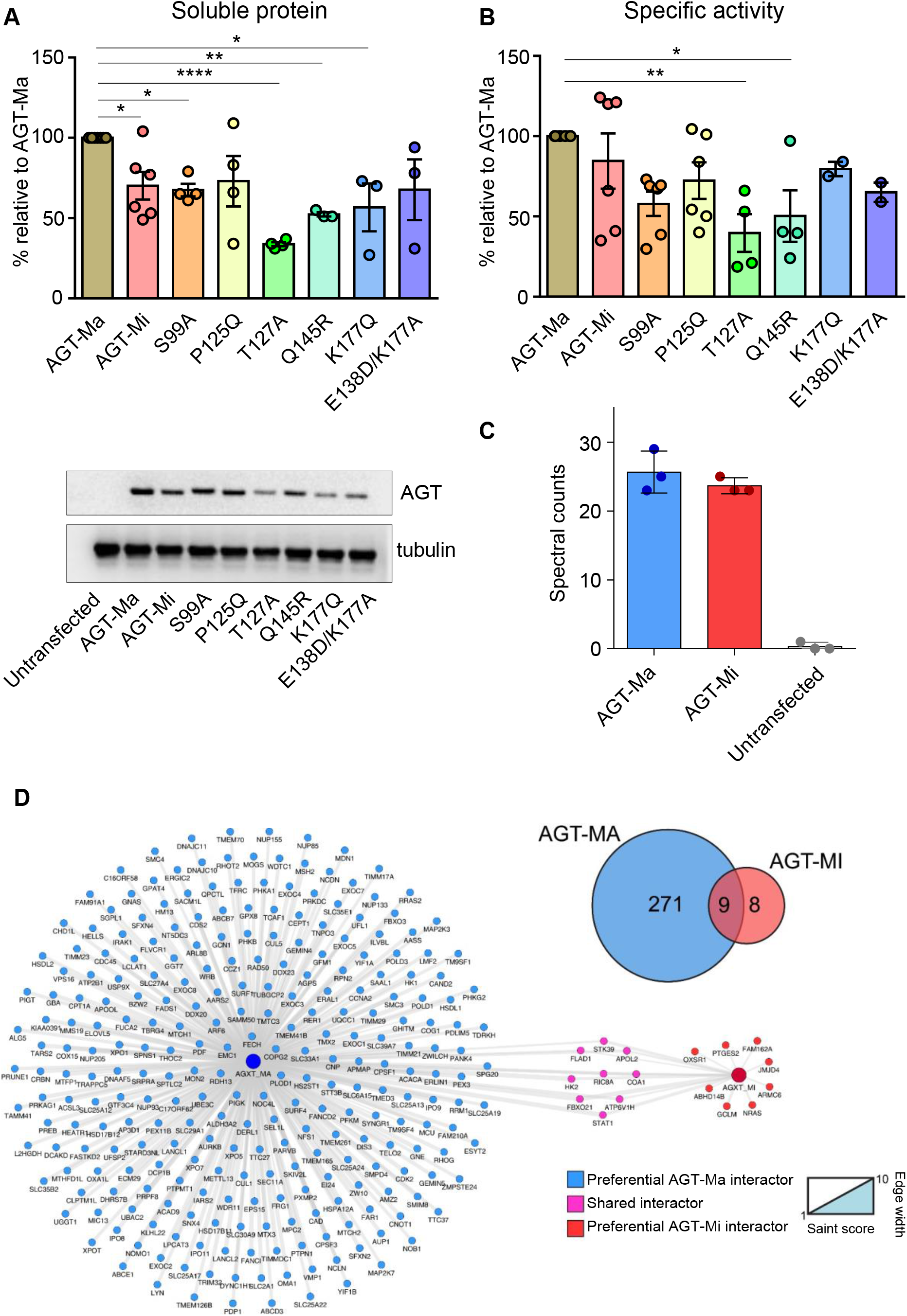
Expression, specific activity and interactors of AGT-Ma, AGT-Mi, and selected variants in human cells. **A) and B)** HEK293 cells expressing the indicated species were harvested and lysed. In panel A, soluble protein levels were quantified by western-blot. 10 μg of soluble lysate of each sample were subjected to SDS-PAGE, immunoblotted with anti-AGT from rabbit (1:10000), and detected by chemiluminescence. The histogram is representative of immunoblot band volume (as mean ± SEM of at least three independent experiments). In panel B, the specific activity for the transamination reaction was measured by incubating 90 μg of soluble lysate with 0.5 M L-alanine, 10 mM glyoxylate, and 200 μM PLP at 25 °C in 100 mM KP, pH 7.4. C-D) Interactome of AGT-Ma and AGT-Mi. Panel C shows that AGT1 protein levels were enriched in AGT-Ma and AGT-Mi transfected samples after immunoprecipitation using a pan anti-AGT1 antibody. Control samples (Ctrl) showed undetectable levels of AGT; Panel D shows Venn diagram of significant interactions compared to controls between AGT-Ma and AGT-Mi and the interaction network of AGT-Ma and AGT-Mi isoforms. Nodes are representative of proteins described with gene name, edges’ width is proportional to SAINT score vs control samples.

As expected, cells expressing AGT-Mi displayed a slightly lower specific activity and protein levels as compared to AGT-Ma. The S99A mutation in IDR1 reduced AGT1 specific activity and protein levels below AGT-Ma. These data are in line with those obtained upon bacterial expression, where the S99A variant showed intermediate properties between AGT-Ma and AGT-Mi when present alone, but all double and triple variants containing the S99A mutation displayed a fitness lower than AGT-Mi. On the other hand, differences between the bacterial and mammalian environment can be observed on variants belonging to the second disordered region. P125Q, T127A and Q145R variants showed specific activity values higher than AGT-Ma upon expression in *E. coli*. However, in analogy with the data obtained on the mutation of His146 (**Fig. S3**), the presence of these mutations strongly impaired the specific activity and relative expression of AGT1 in mammalian cells. Indeed, the P125Q variant showed a fitness comparable to AGT-Mi while a lower fitness is observed in the presence of T127A and Q145R mutations. Since Pro125, Thr127 and Gln145 do not affect the kinetic properties of AGT or its intrinsic thermodynamic stability, these results clearly indicate that specific interactions occurring in the mammalian environment could be responsible for the observed changes. It can be suggested that the conformational change caused by the mutation of Thr127 and Gln145 could favour the interaction with components of the quality control network typical of eukaryotic cells (31). Finally, the mutations in the third disordered region are expected to slightly affect protein fitness with L172A and C173A displaying a specific activity similar, if not higher, compared to AGT-Ma (**Fig. S3**). However, the K177Q and E138D/K177Q variants showed specific activity and proteins levels lower than AGT-Ma and similar to those of AGT-Mi, in line with the observed changes of melting temperature caused by the Lys177 mutation. Overall, these data allow to conclude that introducing mutations in residues belonging to IDRs can affect protein fitness in a cellular environment, suggesting that the potential involvement of IDRs in AGT1 interactions with binding partners.

To analyse the interactome of the two AGT isoforms we used co-immunoprecipitation coupled with quantitative mass spectrometry (IP-MS). HEK293 cells expressing AGT-Ma and AGT-Mi showed a similar number of identified proteins compared to controls not expressing AGT (~3000, data not shown). Label-free quantification of AGT showed a significant enrichment of both AGT-Ma and AGT-Mi isoforms compared to control samples in which AGT was basically not detectable (**Fig. 4C**). SAINT analysis of the interactions of AGT-Ma and AGT-Mi indicated a marked difference in the preferential interactome of the two isoforms (**Table S6**). According to our filtering criteria versus control IPs, AGT-Ma showed 280 significant interactions while AGT-Mi only 17 (**Fig. 4D**). Overlap analysis of the interactions between the two isoforms showed also a minimal overlap with 9 proteins interacting with both AGT-Ma and AGT-Mi.

## DISCUSSION

The main finding of this work is that AGT1 is endowed with a high conformational plasticity (fuzziness), due to the presence of structurally frustrated regions. This leads to two main questions: *how AGT1 structural frustration has been shaped during evolution? Which are the implications of AGT1 instability on the fitness and function at the cellular level?*

The presence of frustrated regions that render an enzyme less tolerant to mutations seems counterintuitive, but AGT1 can be considered a paradigmatic example of how unstable regions are conserved since they possibly mediate protein-protein interactions. In the subtle trade-off between a stable structure, favouring the catalytic activity, and a more flexible one, that may allow multiple interactions, nature finds a compromise features by keeping the degree of disorder within a strict fitness window (**Fig. 5**).

**Fig. 5.**
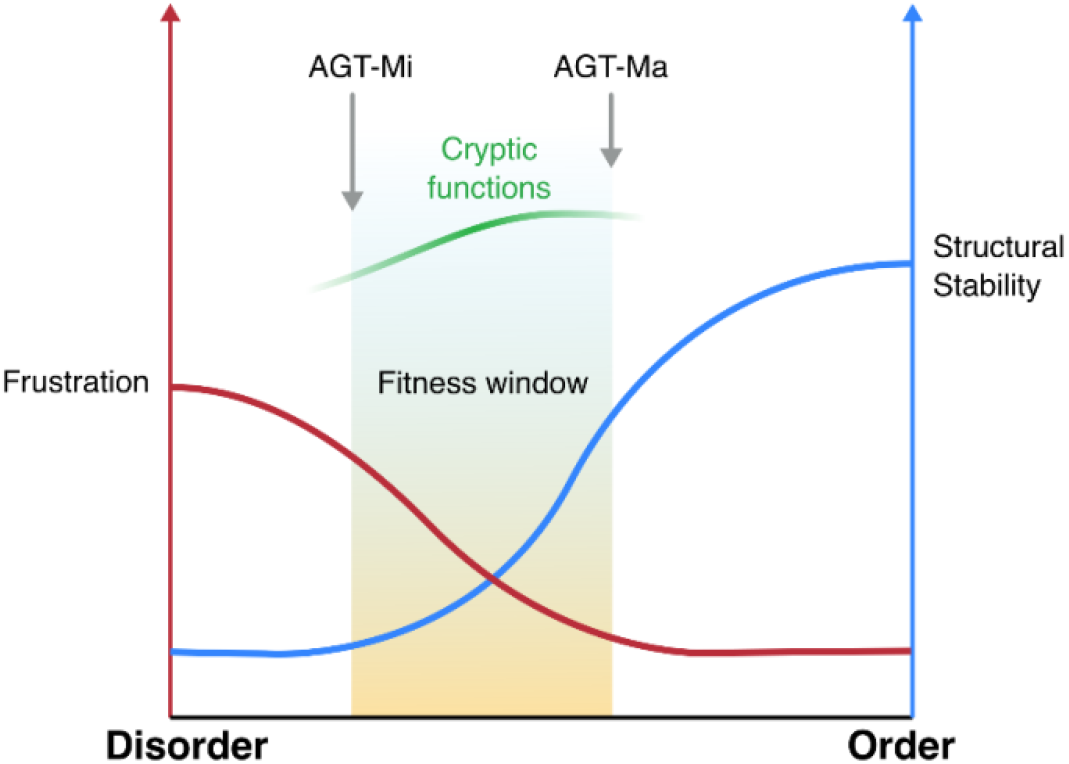
Scheme representing the fitness window of AGT1 as resulting from the order/disorder switch that balances structural stability (blue curve) and frustration (red curve).

As such, we immediately recognise the AGT1 polymorphism as a powerful model system to explore how point mutations may affect protein fitness. AGT-Ma and AGT-Mi structurally differ for the degree of disorder of three peripheral regions, and represent the limits of the functional-fitness window of AGT1 (15). We explored these limits by constructing a small-scale “anti-consensus” library and found that it not only sampled the entire fitness range from AGT-Mi to AGT-Ma, but also identified variants with higher activity and/or stability than AGT-Ma. Although it is not possible to rationalize the effect of each mutation, we noticed, as expected, that mutating less frequent residues to the most frequent ones in other species, give rise to advantageous effects, while the mutation of most conserved residues to less conserved seems to be generally disadvantageous. Since we were able to sample many AGT1 variants with a stability and activity comparable or even better then AGT-Ma with just a small library, it is also surprising that evolution conserves the major allele as it is, without improving its structural stability. This cannot be the outcome of chance and must rather be the result of a necessity. It is likely that the conformational plasticity of AGT-Ma due to the presence of fuzzy regions may have been conserved by evolution because of the balance between the two opposite forces of structural stability and innovability. This would agree with recent studies indicating that proteins showing IDRs play a major role during the first stages of speciation due to their tolerability to mutations (32). Therefore, the evolutionary advantage goes beyond the catalytic efficiency to encompass the stability/function/interactions in a cellular environment. From the structural perspective, perturbation of the frustrated regions with naturally occurring substitutions results in subtle thermostability and activity changes of the purified proteins or in a bacterial background, whereas when the variants are investigated in a eukaryotic cellular context the role of these IDRs is uncovered.

The frustrated regions therefore play a crucial role for AGT stability in the cell with AGT-Ma maximising both the interactions and the activity and AGT-Mi being at the edge of misfolding. Following this view, if the protein is forced outside this window either way (i.e. toward disorder or toward a too stable structure) it can lose one of the functions.

Our data on the behaviour of selected single variants of the library expressed in HEK293 cells agree with this hypothesis, by showing that mutations affecting the frustrated regions that render AGT1 more stable and active when expressed in bacteria (i.e. T127A and Q145R), reduce the intracellular stability in human cells in terms of both protein levels and specific activity. On the other hand, mutations that shift the equilibrium of the native conformers towards a disordered state may induce aggregation or degradation upon sensing by the quality control systems (31). Indeed, the majority of protein stretches that confer protein instability, called degrons, are located in disordered surface regions that facilitate local unfolding and binding to the proteasome (33). Degrons -namely portions of protein that regulate degradation rate- in frustrated regions affect protein half-life and can represent hubs for protein-protein interactions, so that their alterations also lead to interactome rewiring (33, 34). This observation is in line with our proteomic data showing that AGT-Ma is able to establish protein-protein interactions, whose number is remarkably reduced in AGT-Mi, the form mainly shifted toward disorder. Notably, a common binding partner between the two forms is FBXO21, a E3 ubiquitin ligase (35), thus further reinforcing the view that frustrated regions in AGT1 play a crucial role in regulating protein turnover.

Besides structurally uncovering an unexpectedly wide structural landscape of AGT1 in the native state, our data combined with previous evidences also definitely place AGT-Mi as a lower limit of fitness. Interestingly, when looking at the rate of the evolution of the AGT1 structure, positions 11 and 340 are characterised by low evolutionary rate (**Fig. S7**). While Pro11 is well conserved as expected, Ile340 is not conserved and its substitutions to Met represents a consensus change (**Fig. S8** and **S9**) (25, 36). Thus, the P11L is supposed to be highly destabilising and compensated by the I340M change (14). Our structural data allows to elucidate the molecular mechanisms behind the evolutionary changes of AGT1. Indeed, under conditions mimicking a physiological environment in terms of pH, ionic strength and temperature, AGT-Mi in the native form is more prone to aggregation mediated by electrostatic forces (25). Moreover, the two polymorphic mutations P11L and I340M typical of the minor allele oppositely affect the propensity of native AGT1 to aggregation and degradation. Our data confirm and structurally explain these observations as a consequence of the increased surface frustration in AGT-Mi, which is promoted by the P11L mutation at the N-terminus, and attenuated by the I340M mutation, therefore giving a detailed picture of how the two changes compromise to keep the protein in a strict fitness window. Moreover, the data allow to re-interpret the effects of two PH1-associated mutations that occur in region 2 (D129H and E141D) that were predicted to cause the AGT1 deficit by changing the electrostatic potential surface (25). It is likely that these mutations modify the degree of disorder of region 2 with consequent negative impact on the intracellular stability of the protein.

The latter observations call for a third and last question: *why is the minor allele polymorphism retained during evolution?*

Although the detailed analysis of the interactors at functional level goes beyond the scope of this work, we observed that AGT1 seems to retain the possibility to interact with partners involved in mitochondrial protein import and/or assembly (**Table S6**). It cannot be excluded that this property represents a vestigial of the mitochondrial localization of AGT1 that has been lost as a consequence of an adaptive shift in AGT targeting, under the evolutive pressure of a diet change from mainly carnivorous to omnivorous and herbivorous (37). Notably, the P11L polymorphism on the one hand reduces protein half-time and remodels regions possibly involved in the interaction with binding partners, but also generates a weak N-terminal mitochondrial targeting sequence that redirects 5% of the protein to mitochondria, where the metabolism of hydroxyproline takes place (14). It can be speculated that this feature might be conserved in Caucasian population, whose diet includes more meat than African and Asian populations, as an advantage which may in part compensate for the reduced activity and interactions. This view opens new avenues for studies aimed at better understanding glyoxylate/oxalate metabolism in humans and its relation with environmental conditions.

Overall, our work has broader implications into our view of proteins that display a flat-bottom folding funnel, because the native state is not unique but fluctuates among different conformers. The conformational landscape explored by such proteins can be greatly influenced by either environmental changes or naturally-occurring mutations. Following this idea, and especially in the case of metabolic enzymes, the fitness in a whole cell depends on a pool of different conformers, and it is linked to the necessity of balancing catalysis with networking. From an evolutionary point of view, the fate of a protein would then not only depend on the generation of partially-folded intermediates, but also from the equilibrium between “functional” or “potentially noxious” native conformers, which in turn might be determined by surface residues whose importance has been underestimated in classical biochemical and protein folding studies. In this regard, a growing number of reports underline that natural polymorphisms, including those related to IDRs, as key contributors to human diseases (38, 39), and that single amino acid changes common in human population more likely affect binding interfaces with other proteins or nucleic acids (40).

The AGT1 case can thus be viewed as an example of evolutionary “risk-management”: a certain degree of instability and disorder, although it may lead to misfunction, is retained and “controlled” in order to achieve additional or cryptic functions (multitasking).

## Materials and Methods

*Reagents-*Pyridoxal phosphate (PLP), L-alanine, sodium glyoxylate, rabbit muscle L-lactic dehydrogenase (LDH), β-Nicotinamide adenine dinucleotide reduced form (NADH) and isopropyl-β-D-thiogalactoside (IPTG) were purchased from Sigma-Aldrich. Protease Inhibitor Cocktails organic solvents were purchased from Nacalai tesque. All other chemicals unless otherwise noted, were from Sigma-Aldrich, Nacalai tesque or Wako. The anti-AGT1 antibody from rabbit was provided by Prof. Chris Danpure (University College London).

*Crystallography of AGT-Mi and structure determination-*Single crystals of AGT-Mi were obtained by vapor diffusion methods (sitting drop) mixing 0.4 μl of a 37 μM protein solution (buffer: 0.1 M potassium phosphate pH=7.4) with 0.4 μl of the A9 solution of the Morpheus screen (Molecular Dimension), containing: 30 mM MgCl 2, 30 mM CaCl 2, 50 mM sodium 4-(2-hydroxyethyl)-1-piperazineethanesulfonate (HEPES) + 50 mM 3-(N-morpholino) propane sulfonic acid (MOPS) at a final pH 7.5, 40% v/v Ethylene glycol; 20 % w/v PEG 8K. Yellow diamond-shaped crystals of about 100-120 μm grew in 1 week. Crystals were flash frozen without prior addition of cryoprotectant. A complete X-ray diffraction dataset was collected at the Elettra synchrotron in Trieste (IT) at XRD1 beamline. The data were processed with XDS 2 and scaled with Aimless 3–5 initially in P4_1_ 2_1_ 2 space group to a final resolution of 2.0 Å. However, the observed intensities were very different from the theorical values suggesting the presence of twinning. Indeed, although a very clear molecular replacement solution was found in P4_1_ 2_1_ 2 - using the AGT-Ma structure as search model (pdb: 5F9S6) - and the electron density maps were very clear, the model refined to poor R free and geometry values with respect to the resolution of the data. Therefore, different space groups were evaluated and finally the data were reindexed in P4_1_ with merohedral twin operator h, −k, −l and an estimated twin fraction of 0.44 (Phenix.xtriage 7) that converged to 0.49 during twin refinement in Refmac5 or Phenix.refine 7–9. Given the almost perfect twin, model building was carried out in Coot 10 using electron density maps calculated ignoring twinning, since twin-refinement may introduce a strong model-bias, and with an extensive and careful use of omit-maps. Twin-refinement was used only in the final refinement cycle. Data collection and refinement statistics are summarized in **Table S1** and the quality of the electron density maps is illustrated in **Fig. S1**. Coordinates and structure factors have been deposited in the protein data bank with pdb code: 7NS7. B-factor analysis of AGT-Ma solved structures was performed in Phenix GUI using structure comparison tool 7. Figures were rendered using Chimera 11.

*Molecular Dynamics of AGT-Mi and AGT-Ma-*The dimeric structure of AGT-Ma (PDB ID 5F9S) was used as starting point of MD simulations, and as template for the generation of the mutant structures. The monomeric structure of the mutants was obtained through the Swiss-model web server (41), realigned to the reference structure, and manually added of the PLP atoms. PDB file manipulation was performed using pdb-tools (42). By following the procedure of Gupta *et al.,* a hybrid residue named ‘LYP’ was created by combining the PLP and the Lysine 209 atoms in the PDB file and manually parametrized in the forcefield using the same values reported in the paper (43). All simulations were performed using GROMACS version 2020.3 (www.gromacs.org) (44) and CHARMM36-mar2019 forcefield (45). The model was solvated in a dodecahedral box using 3 points charge water model (tip3p) (46). After charge, temperature, and pressure equilibration, each simulation run 200 ns in 2 fs steps (four repeats). A modified Berendsen thermostat was used to maintain the temperature at 310K (47), and a Parrinello-Rahman barostat was used to maintain a pressure of 1 bar (48). A Particle Mesh Ewald (PME) was used for long-range electrostatic interactions (49). More details about the system setup and simulation parameters are provided as Supplementary Information.

*Expression and purification of AGT1 and variants-E. coli* BL21(DE3) cells were transformed with the constructs encoding AGT-Ma, AGT-Mi and the selected variants of the library. Expression and purification of the proteins were performed as reported (9).

*Site directed mutagenesis-*Mutants were constructed as described in the Q5 Site-Directed Mutagenesis manual (New England Biolabs). The primers list is reported in **Table S4**.

*Frustration analysis-*Analysis of protein frustration was performed by uploading the AGT-Ma structure (pdb: 5F9S) (50) to the Frustratometer server (http://frustratometer.qb.fcen.uba.ar/) (27) First a list of interacting residues are computed (Q_i,j_) and the native energy of the structure is calculated according to the AMW energy function (51). Frustration between a given couple of amino acids (i,j) is then calculated by perturbating the protein sequence and calculating the total energy change. The contact is then considered minimally frustrated, neutral or highly frustrated according to how favourable the interaction is compared to all possible interaction. Conformational frustration was visualized using VMD molecular graphics software (52).

*Kinetics Assays-*To determine the kinetic parameters for the transamination reaction, the purified proteins (0.1-0.2 μM) were incubated in presence of 100 mM PLP in KP 0.1 M pH 7.4 at 25 °C at increasing alanine concentrations (15 to 500 mM) or glyoxylate concentrations (0.07 to 5 mM) and a fixed glyoxylate (10 mM) or alanine (500 mM) concentration. The reaction was quenched by adding TCA 10% (v/v) and pyruvate production was measured using a spectrophotometric assay coupled with LDH following the NADH signal at 340 nm (9). To determine the activity in crude bacterial cell lysates, a Tecan Spark Microplate Reader (Thermo Fischer Scientific) was used and the cells were grown and lysed as described below under the section ‘Library Screening’.

To measure the enzymatic activity in HEK293 cellular lysates, 100 μg of lysate were incubated with 0.5 M L-alanine and 10 mM glyoxylate at 25°C for 20-60 min in 100 mM KP buffer, pH 7.4 in presence of 100 μM PLP. The reactions were stopped by adding TCA 10% (v/v) and pyruvate production was measured using a spectrophotometric assay coupled with lactate dehydrogenase (53).

*Circular Dichroism spectroscopy*- Circular Dichroism (CD) spectra were obtained using a Jasco J-820 spectropolarimeter with a thermostatically controlled compartment at 25 °C. In the near-UV and visible spectra, protein concentration was 10 μM in a cuvette with a path length of 1 cm. Far-UV measurements were taken at a protein concentration of 0.1 mg/ml using a cuvette with a path length of 0.1 cm. The thermal unfolding of AGT-Ma, AGT-Mi and variants was monitored by following the loss of CD signal at 222 nm at a protein concentration of 5-8 μM with temperature increasing of 0.5 °C/min from 25 to 90 °C. The experiments were performed in KP 0.1 M pH 7.4.

*Cloning and library construction-*The AGT1 sequence (human) (uniprot: P21549, SPYA_HUMAN) was used for conducting NCBI BLAST searches across the non-redundant database. Database search conditions were filtered for search results with query coverage >80% and sequence identity between 70-90% were considered for the next step. The collected sequences were then subjected to multiple sequence alignment using MUltiple Sequence Comparison by Log-Expectation (MUSCLE). The library was prepared using a modified Combinatorial Assembly PCR protocol. The three different regions of interest (97-102, 121-146 and 170-177) were mutated using single stranded mutagenic degenerated primers (**Table S5**) that anneal each other at their edges. The PCR fragments of the three regions of interest were cloned in the pTrcHis2A plasmid by Megawhop cloning (54) and mixed between them to obtain random combination of double and triple mutants. The library was then transformed in BL21 (DE3) *E. coli* strain and more than 2000 clones were screened.

*Library screening-*Library has been cloned into pTrcHis-2A, transformed into *E. coli* BL21 (DE3) and plated on ampicillin LB-agar plates. Colonies were picked into deep 96-well plates and grown overnight at 30 °C in 500 μL LB per well supplemented with 50 μg/mL ampicillin. Wells containing fresh LB/ampicillin (2 mL) were inoculated with 100 μL of this preculture, and protein expression was induced adding 0.1 mM IPTG at 25 °C for 16-18 hours. Cells were pelleted by centrifugation and the supernatant removed. Cell lysis was performed by freezing for 60 minutes at −80 °C, and resuspending in 200 μL of sodium phosphate buffer pH 7.4 and supplemented with 100 μg/mL lysozyme, proteinase inhibitor cocktail (Nacalai tesque) and 25 U/mL benzonase (Merck). The mixture has been incubated for 30 min at room temperature. Cell debris was pelleted by centrifugation and 20–100 μL of clarified lysate were removed and tested for AGT1 activity. The activity was tested in presence of 10 μM PLP and saturating concentration of substrate and co-substrate (10 mM glyoxylate and 500 mM l-alanine) in 200 μL of KP 0.1 M pH 7.4. Some variants were picked and regrown in triplicate, and the rates re-measured and the average values determined. Plasmid DNA was isolated, and variants were sequenced. The plasmids were retransformed, and the resulting chosen mutants were used to inoculate new cultures for protein purification.

*Cell culture, transfection, and lysis-*Human Embryonic Kidney cells (HEK293 cells) were cultured in Dulbecco’s modified Eagle’s medium (DMEM) supplemented with fetal bovine serum (FBS) (10%, v/v), penicillin (100 units/ml) and streptomycin (100 μg/ml) at 37 °C in a 5% CO_2_ humidified environment. To test the effects of different AGT1 single amino-acid substitutions on the expression level and enzymatic activity of AGT, cells were seeded in Petri dishes at the density of 2.1×10^4^/cm^2^ in DMEM. After 24h of subculture, cells were transfected with different AGT1 constructs using Lipofectamine LTX and PLUS Reagent (Invitrogen) according to the manufacturer’s instructions. After 24 h, cells were harvested and used for Western Blotting analyses and enzymatic activity assays. For Co-IP experiments, stable clones of HEK293 cells expressing AGT-Ma or AGT-Mi were generated by selecting transfected cells with 1 mg/ml G418 in the culture medium for three weeks.

For enzymatic activity assays and western-blot analyses cell pellets were lysed by freeze/thawing (five cycles) in PBS (Phosphate Buffer Saline), pH 7.2 supplemented with 100 μM PLP and protease inhibitor cocktail (CompleteMini, Roche). The whole cell extract was separated by centrifugation (12,900 x *g,* 10 min, 4 °C) to obtain the soluble fraction. Protein concentration in the soluble fraction was measured using the Bradford protein assay.

*Western-Blot and Co-immunoprecipitation-*10 μg of soluble cell lysate were loaded on 10% SDS-PAGE, transferred on a nitrocellulose membrane and immunoblotted with anti AGT antibody (1:10.000) in 2.5% milk w/v in TTBS (50mM Tris-HCl, pH 7.5, 150 mM NaCl, 0.1% Tween 20) overnight at 4°C. After three washes in TBST, the membrane was incubated with peroxidase-conjugated antirabbit immunoglobulin G (IgG) (1:10.000) in 5% milk in TTBS for 1 hour at room temperature. Immunocomplexes were visualized by an enhanced chemiluminescence kit (ECL, Pierce Biotechnology, Rockford, IL). An antibody against human tubulin was used as internal loading control.

Co-immunoprecipitation was performed using the Pierce™ Co-Immunoprecipitation Kit according to manufacturer’s instructions (Thermo-Scientific). Briefly, cell pellets were lysed in the IP Lyses/Wash Buffer included in the kit, supplemented with protease inhibitor cocktail (CompleteMini, Roche) and phosphatase inhibitors (Sigma-Aldrich). The whole cell extract was separated by centrifugation (12,900 x *g,* 10 min, 4 °C) to obtain the soluble fraction. Two mg of soluble cell lysate from untransfected (negative control) or transfected cells (AGT-Ma and AGT-Mi) were incubated with 10 μg of molyclonal anti-AGT primary antibody, immobilized on column according to the manufacturer’s instructions, overnight at 4° C. The samples were eluted from the column using SDS buffer (2% SDS, 100 mM Tris-HCl pH 7.2, 10 mM DTT). After immunoprecipitation, the protein eluates were precipitated overnight with ice-cold acetone (1:4 v/v) and subjected to sample preparation for proteomics according to previously published procedures (55).

*Mass-spectrometry and data analysis-*Peptides derived from tryptic digestion (~1 μg) were separated on a reverse phase PicoFrit column (75 um ID, 8 Um tip, 250 mm bed packed with Reprosil-PUR C18-AQ, 1.9 μm particle size, 120 Å pore size, New Objective, Inc., Woburn, MA, USA, cat. PF7508-250H363), using an EASY-nLC™ 1200 System (Thermo Fisher Scientific, Waltham, MA, USA). Total run time for each sample was 120 min, peptides were separated using a 100 min gradient (4–40% acetonitrile +0.1% formic acid at 300 nL/min). Eluting peptides were measured on-line with a Q Exactive HF benchtop mass spectrometer (Thermo Fisher Scientific) operating in data-dependent acquisition mode (Top20). Peptides were ionized at a potential of +2 KV, intact peptide ions were detected at a resolution of 120,000 (at m/z 200) and fragment ions at a resolution of 15,000 (at m/z 200); AGC Target settings for MS were 3·10^6^ charges and for MS/MS 1·10^5^ charges. Peptides were selected for Higher-energy C-trap dissociation (HCD) fragmentation with a quadrupole isolation window of 1.4 Th, peptides were fragmented at a normalized collision energy of 30. The intensity threshold was set at 2·10^4^ and Dynamic exclusion at 30 s. Raw files from MS analysis were processed using the MaxQuant software version 1.6.10.43 22 (Martinsried, Germany) [15]. The spectra were searched against a Uniprot human database (release 2020_2, 20,375 sequences, including isoforms). Precursor and fragment mass tolerance were set to 7 and 20 ppm., respectively, while the minimum length of the peptide was set to 7. False discovery rate (FDR) was set to 1% both at the peptide and protein level. Enzyme for *in silico* digestion were set to trypsin and lysine C, and up to two missed cleavages were allowed. Cysteine carbamidomethylation (Cys, +57.021464 Da) was set as a fixed modification, whereas N-acetylation of proteins (N-terminal, +42.010565 Da) and oxidized methionine (Met, +15.994915 Da) were included as variable modifications.

Proteins identified and quantified via LC-MS/MS were analysed using different bioinformatics tools to distinguish interactors from false positives and obtain a list of candidates constituting the core local network of AGT proteoforms. Label-free quantification of proteins were performed using spectral counts, while the Significance Analysis of INTeractome (SAINT) approach was used for scoring the significance of the interactions (56–58). This method estimates the distribution of protein abundances under true and false interaction hypotheses and computes the probability of true interaction, taking into account the information from negative controls included in the experimental setup. Our negative controls were represented by wild-type HEK293 cells not expressing AGT (Ctrl, n=3) and HEK293 expressing the AGT-Ma or AGT-Mi isoforms subjected to the co-IP without the AGT antibody. Scored interaction by the SAINT algorithm were filtered using a fold change vs controls > 2 and a SAINT score (SP score)>0.7.

Network visualization was performed using the Cytoscape software as previously reported (59). The mass spectrometry proteomics data have been deposited to the ProteomeXchange Consortium via the PRIDE (60) partner repository with the dataset identifier PXD025661.

### Phylogenetic analysis

The aligned sequences of 254 annotated vertebrate AGT1 was obtained from the Ensembl database (61). The best-fit evolutionary model (JTT+6R) (62, 63) was chosen using ModelFinder (64). The phylogenetic tree and the rates of evolution were calculated using IQTREE software (65). The Coelacanth sequence ENSLACT00000026005 was used as outgroup for the tree generation.

## Author contributions

B.C., G.G., P.L. and M.D. conceived the project. M.D. and P.L. planned the library. M.D. and G.U. generated the library and tested the activity. M.D. expressed, purified and characterised single mutants. S.P. performed molecular dynamics simulations, rate of evolution and phylogenetic analysis. G.G. collected crystallography data, processed, solved and analysed crystallography data. S.G. and D.C. performed experiments in Hek293 cells and analysed MS data. C.C. analyzed Hek293 and MS data and revised the manuscript. B.C., P.L. and G.G. supervised the project and wrote the manuscript with the input from all the authors.

## Acknowledgments

The authors would like to thank the Scientific Computing and Data Analysis section of Research Support Division at OIST for the support provided for the molecular dynamic simulations. The authors would also like to thank Stefano Gianni and Benjamin Clifton for critical reading and feedback on the manuscript.

## References

1 Gibson, G. and Dworkin, I. (2004) Uncovering cryptic genetic variation. Nature reviews. Genetics, 5, 681–690.

2 Paaby, A.B. and Rockman, M.V. (2014) Cryptic genetic variation: evolution’s hidden substrate. Nature reviews. Genetics, 15, 247–258.

3 Zheng, J., Payne, J.L. and Wagner, A. (2019) Cryptic genetic variation accelerates evolution by opening access to diverse adaptive peaks. Science, 365, 347–353.

4 Baier, F., Hong, N., Yang, G., Pabis, A., Miton, C.M., Barrozo, A., Carr, P.D., Kamerlin, S.C., Jackson, C.J. and Tokuriki, N. (2019) Cryptic genetic variation shapes the adaptive evolutionary potential of enzymes. Elife, 8.

5 Gianni, S., Freiberger, M.I., Jemth, P., Ferreiro, D.U., Wolynes, P.G. and Fuxreiter, M. (2021) Fuzziness and Frustration in the Energy Landscape of Protein Folding, Function, and Assembly. Acc Chem Res, 54, 1251–1259.

6 Uversky, V.N. (2019) Intrinsically Disordered Proteins and Their “Mysterious” (Meta)Physics. Frontiers in Physics, 7.

7 Babu, M.M. (2016) The contribution of intrinsically disordered regions to protein function, cellular complexity, and human disease. Biochemical Society transactions, 44, 1185–1200.

8 Liu, Z. and Huang, Y. (2014) Advantages of proteins being disordered. Protein science: a publication of the Protein Society, 23, 539–550.

9 Cellini, B., Bertoldi, M., Montioli, R., Paiardini, A. and Borri Voltattorni, C. (2007) Human wild-type alanine:glyoxylate aminotransferase and its naturally occurring G82E variant: functional properties and physiological implications. The Biochemical journal, 408, 39–50.

10 Oppici, E., Montioli, R. and Cellini, B. (2015) Liver peroxisomal alanine:glyoxylate aminotransferase and the effects of mutations associated with Primary Hyperoxaluria Type I: An overview. Biochimica et biophysica acta, 1854, 1212–1219.

11 Lorenzo, V., Torres, A. and Salido, E. (2014) Primary hyperoxaluria. Nefrologia: publicacion oficial de la Sociedad Espanola Nefrologia, 34, 398–412.

12 Purdue, P.E., Takada, Y. and Danpure, C.J. (1990) Identification of mutations associated with peroxisome-to-mitochondrion mistargeting of alanine/glyoxylate aminotransferase in primary hyperoxaluria type 1. The Journal of cell biology, 111, 2341–2351.

13 Lumb, M.J. and Danpure, C.J. (2000) Functional synergism between the most common polymorphism in human alanine:glyoxylate aminotransferase and four of the most common disease-causing mutations. The Journal of biological chemistry, 275, 36415–36422.

14 Danpure, C.J. (2006) Primary hyperoxaluria type 1: AGT mistargeting highlights the fundamental differences between the peroxisomal and mitochondrial protein import pathways. Biochimica et biophysica acta, 1763, 1776–1784.

15 Mesa-Torres, N., Salido, E. and Pey, A.L. (2014) The lower limits for protein stability and foldability in primary hyperoxaluria type I. Biochimica et biophysica acta, 1844, 2355–2365.

16 Cellini, B., Montioli, R. and Voltattorni, C.B. (2011) Human liver peroxisomal alanine:glyoxylate aminotransferase: characterization of the two allelic forms and their pathogenic variants. Biochimica et biophysica acta, 1814, 1577–1584.

17 Hopper, E.D., Pittman, A.M., Fitzgerald, M.C. and Tucker, C.L. (2008) In vivo and in vitro examination of stability of primary hyperoxaluria-associated human alanine:glyoxylate aminotransferase. The Journal of biological chemistry, 283, 30493–30502.

18 Mesa-Torres, N., Fabelo-Rosa, I., Riverol, D., Yunta, C., Albert, A., Salido, E. and Pey, A.L. (2013) The role of protein denaturation energetics and molecular chaperones in the aggregation and mistargeting of mutants causing primary hyperoxaluria type I. PloS one, 8, e71963.

19 Cellini, B., Lorenzetto, A., Montioli, R., Oppici, E. and Voltattorni, C.B. (2010) Human liver peroxisomal alanine:glyoxylate aminotransferase: Different stability under chemical stress of the major allele, the minor allele, and its pathogenic G170R variant. Biochimie, 92, 1801–1811.

20 Montioli, R., Fargue, S., Lewin, J., Zamparelli, C., Danpure, C.J., Borri Voltattorni, C. and Cellini, B. (2012) The N-terminal extension is essential for the formation of the active dimeric structure of liver peroxisomal alanine:glyoxylate aminotransferase. The international journal of biochemistry & cell biology, 44, 536–546.

21 Dindo, M., Mandrile, G., Conter, C., Montone, R., Giachino, D., Pelle, A., Costantini, C. and Cellini, B. (2020) The ILE56 mutation on different genetic backgrounds of alanine:glyoxylate aminotransferase: Clinical features and biochemical characterization. Molecular genetics and metabolism, 131, 171–180.

22 Cellini, B., Montioli, R., Paiardini, A., Lorenzetto, A., Maset, F., Bellini, T., Oppici, E. and Voltattorni, C.B. (2010) Molecular defects of the glycine 41 variants of alanine glyoxylate aminotransferase associated with primary hyperoxaluria type I. Proceedings of the National Academy of Sciences of the United States of America, 107, 2896–2901.

23 van der Lee, R., Buljan, M., Lang, B., Weatheritt, R.J., Daughdrill, G.W., Dunker, A.K., Fuxreiter, M., Gough, J., Gsponer, J., Jones, D.T. et al. (2014) Classification of intrinsically disordered regions and proteins. Chem Rev, 114, 6589–6631.

24 Jakob, U., Kriwacki, R. and Uversky, V.N. (2014) Conditionally and transiently disordered proteins: awakening cryptic disorder to regulate protein function. Chem Rev, 114, 6779–6805.

25 Dindo, M., Conter, C. and Cellini, B. (2017) Electrostatic interactions drive native-like aggregation of human alanine:glyoxylate aminostransferase. The FEBS journal, 284, 3739–3764.

26 Goddard, T.D., Huang, C.C., Meng, E.C., Pettersen, E.F., Couch, G.S., Morris, J.H. and Ferrin, T.E. (2018) UCSF ChimeraX: Meeting modern challenges in visualization and analysis. Protein science: a publication of the Protein Society, 27, 14–25.

27 Parra, R.G., Schafer, N.P., Radusky, L.G., Tsai, M.Y., Guzovsky, A.B., Wolynes, P.G. and Ferreiro, D.U. (2016) Protein Frustratometer 2: a tool to localize energetic frustration in protein molecules, now with electrostatics. Nucleic acids research, 44, W356–360.

28 Strodel, B., Whittleston, C.S. and Wales, D.J. (2007) Thermodynamics and kinetics of aggregation for the GNNQQNY peptide. J Am Chem Soc, 129, 16005–16014.

29 Waterhouse, A.M., Procter, J.B., Martin, D.M., Clamp, M. and Barton, G.J. (2009) Jalview Version 2--a multiple sequence alignment editor and analysis workbench. Bioinformatics, 25, 1189–1191.

30 Cellini, B., Montioli, R., Paiardini, A., Lorenzetto, A. and Voltattorni, C.B. (2009) Molecular Insight into the Synergism between the Minor Allele of Human Liver Peroxisomal Alanine:Glyoxylate Aminotransferase and the F152I Mutation. The Journal of biological chemistry, 284, 8349–8358.

31 Fernandez-Higuero, J.A., Betancor-Fernandez, I., Mesa-Torres, N., Muga, A., Salido, E. and Pey, A.L. (2019) Structural and functional insights on the roles of molecular chaperones in the mistargeting and aggregation phenotypes associated with primary hyperoxaluria type I. Adv Protein Chem Struct Biol, 114, 119–152.

32 Forcelloni, S. and Giansanti, A. (2020) Mutations in disordered proteins as early indicators of nucleic acid changes triggering speciation. Scientific reports, 10, 4467.

33 Guharoy, M., Bhowmick, P., Sallam, M. and Tompa, P. (2016) Tripartite degrons confer diversity and specificity on regulated protein degradation in the ubiquitin-proteasome system. Nature communications, 7, 10239.

34 Niemeyer, M., Moreno Castillo, E., Ihling, C.H., Iacobucci, C., Wilde, V., Hellmuth, A., Hoehenwarter, W., Samodelov, S.L., Zurbriggen, M.D., Kastritis, P.L. et al. (2020) Flexibility of intrinsically disordered degrons in AUX/IAA proteins reinforces auxin co-receptor assemblies. Nature communications, 11, 2277.

35 Zhang, C., Li, X., Adelmant, G., Dobbins, J., Geisen, C., Oser, M.G., Wucherpfenning, K.W., Marto, J.A. and Kaelin, W.G., Jr. (2015) Peptidic degron in EID1 is recognized by an SCF E3 ligase complex containing the orphan F-box protein FBXO21. Proceedings of the National Academy of Sciences of the United States of America, 112, 15372–15377.

36 Mesa-Torres, N., Yunta, C., Fabelo-Rosa, I., Gonzalez-Rubio, J.M., Sanchez-Ruiz, J.M., Salido, E., Albert, A. and Pey, A.L. (2014) The consensus-based approach for gene/enzyme replacement therapies and crystallization strategies: the case of human alanine-glyoxylate aminotransferase. The Biochemical journal, 462, 453–463.

37 Birdsey, G.M., Lewin, J., Cunningham, A.A., Bruford, M.W. and Danpure, C.J. (2004) Differential enzyme targeting as an evolutionary adaptation to herbivory in carnivora. Molecular biology and evolution, 21, 632–646.

38 Schuch, J.B., Paixao-Cortes, V.R., Friedrich, D.C. and Tovo-Rodrigues, L. (2016) The contribution of protein intrinsic disorder to understand the role of genetic variants uncovered by autism spectrum disorders exome studies. Am J Med Genet B Neuropsychiatr Genet, 171B, 479–491.

39 Vacic, V. and Iakoucheva, L.M. (2012) Disease mutations in disordered regions-- exception to the rule? Molecular bioSystems, 8, 27–32.

40 Qiu, J., Nechaev, D. and Rost, B. (2020) Protein-protein and protein-nucleic acid binding residues important for common and rare sequence variants in human. BMC Bioinformatics, 21, 452.

41 Waterhouse, A., Bertoni, M., Bienert, S., Studer, G., Tauriello, G., Gumienny, R., Heer, F.T., de Beer, T.A.P., Rempfer, C., Bordoli, L. et al. (2018) SWISS-MODEL: homology modelling of protein structures and complexes. Nucleic acids research, 46, W296–W303.

42 Rodrigues, J., Teixeira, J.M.C., Trellet, M. and Bonvin, A. (2018) pdb-tools: a swiss army knife for molecular structures. F1000Res, 7, 1961.

43 Gupta, S., Kelow, S., Wang, L., Andrake, M.D., Dunbrack, R.L., Jr. and Kruger, W.D. (2018) Mouse modeling and structural analysis of the p.G307S mutation in human cystathionine beta-synthase (CBS) reveal effects on CBS activity but not stability. The Journal of biological chemistry, 293, 13921–13931.

44 Abraham, M.J., Murtola, T., Schulz, R., Páll, S., Smith, J.C., Hess, B. and Lindahl, E. (2015) GROMACS: High performance molecular simulations through multi-level parallelism from laptops to supercomputers. SoftwareX, 1-2, 19–25.

45 Brooks, B.R., Brooks, C.L., 3rd, Mackerell, A.D., Jr., Nilsson, L., Petrella, R.J., Roux, B., Won, Y., Archontis, G., Bartels, C., Boresch, S. et al. (2009) CHARMM: the biomolecular simulation program. J Comput Chem, 30, 1545–1614.

46 Izadi, S., Anandakrishnan, R. and Onufriev, A.V. (2014) Building Water Models: A Different Approach. J Phys Chem Lett, 5, 3863–3871.

47 Bussi, G., Donadio, D. and Parrinello, M. (2007) Canonical sampling through velocity rescaling. J Chem Phys, 126, 014101.

48 Parrinello, M. and Rahman, A. (1981) Polymorphic transitions in single crystals: A new molecular dynamics method. Journal of Applied Physics, 52, 7182–7190.

49 Darden, T., York, D. and Pedersen, L. (1993) Particle mesh Ewald: An N⋅log(N) method for Ewald sums in large systems. The Journal of Chemical Physics, 98, 10089–10092.

50 Giardina, G., Paiardini, A., Montioli, R., Cellini, B., Voltattorni, C.B. and Cutruzzola, F. (2017) Radiation damage at the active site of human alanine:glyoxylate aminotransferase reveals that the cofactor position is finely tuned during catalysis. Scientific reports, 7, 11704.

51 Papoian, G.A., Ulander, J., Eastwood, M.P., Luthey-Schulten, Z. and Wolynes, P.G. (2004) Water in protein structure prediction. Proceedings of the National Academy of Sciences of the United States of America, 101, 3352–3357.

52 Humphrey, W., Dalke, A. and Schulten, K. (1996) VMD: visual molecular dynamics. J Mol Graph, 14, 33–38, 27-38.

53 Oppici, E., Roncador, A., Montioli, R., Bianconi, S. and Cellini, B. (2013) Gly161 mutations associated with Primary Hyperoxaluria Type I induce the cytosolic aggregation and the intracellular degradation of the apo-form of alanine:glyoxylate aminotransferase. Biochimica et biophysica acta, 1832, 2277–2288.

54 Miyazaki, K. (2011) MEGAWHOP cloning: a method of creating random mutagenesis libraries via megaprimer PCR of whole plasmids. Methods in enzymology, 498, 399–406.

55 Macchioni, L., Chiasserini, D., Mezzasoma, L., Davidescu, M., Orvietani, P.L., Fettucciari, K., Salviati, L., Cellini, B. and Bellezza, I. (2020) Crosstalk between Long-Term Sublethal Oxidative Stress and Detrimental Inflammation as Potential Drivers for Age-Related Retinal Degeneration. Antioxidants (Basel), 10.

56 Choi, H., Larsen, B., Lin, Z.Y., Breitkreutz, A., Mellacheruvu, D., Fermin, D., Qin, Z.S., Tyers, M., Gingras, A.C. and Nesvizhskii, A.I. (2011) SAINT: probabilistic scoring of affinity purification-mass spectrometry data. Nat Methods, 8, 70–73.

57 Choi, H., Glatter, T., Gstaiger, M. and Nesvizhskii, A.I. (2012) SAINT-MS1: protein-protein interaction scoring using label-free intensity data in affinity purification-mass spectrometry experiments. J Proteome Res, 11, 2619–2624.

58 Guard, S.E., Ebmeier, C.C. and Old, W.M. (2019) Label-Free Immunoprecipitation Mass Spectrometry Workflow for Large-scale Nuclear Interactome Profiling. J Vis Exp.

59 Chiasserini, D., van Weering, J.R., Piersma, S.R., Pham, T.V., Malekzadeh, A., Teunissen, C.E., de Wit, H. and Jimenez, C.R. (2014) Proteomic analysis of cerebrospinal fluid extracellular vesicles: a comprehensive dataset. J Proteomics, 106, 191–204.

60 Perez-Riverol, Y., Csordas, A., Bai, J., Bernal-Llinares, M., Hewapathirana, S., Kundu, D.J., Inuganti, A., Griss, J., Mayer, G., Eisenacher, M. et al. (2019) The PRIDE database and related tools and resources in 2019: improving support for quantification data. Nucleic acids research, 47, D442–D450.

61 Zerbino, D.R., Achuthan, P., Akanni, W., Amode, M.R., Barrell, D., Bhai, J., Billis, K., Cummins, C., Gall, A., Giron, C.G. et al. (2018) Ensembl 2018. Nucleic acids research, 46, D754–D761.

62 Jones, D.T., Taylor, W.R. and Thornton, J.M. (1992) The rapid generation of mutation data matrices from protein sequences. Comput Appl Biosci, 8, 275–282.

63 Soubrier, J., Steel, M., Lee, M.S., Der Sarkissian, C., Guindon, S., Ho, S.Y. and Cooper, A. (2012) The influence of rate heterogeneity among sites on the time dependence of molecular rates. Molecular biology and evolution, 29, 3345–3358.

64 Kalyaanamoorthy, S., Minh, B.Q., Wong, T.K.F., von Haeseler, A. and Jermiin, L.S. (2017) ModelFinder: fast model selection for accurate phylogenetic estimates. Nat Methods, 14, 587–589.

65 Nguyen, L.T., Schmidt, H.A., von Haeseler, A. and Minh, B.Q. (2015) IQ-TREE: a fast and effective stochastic algorithm for estimating maximum-likelihood phylogenies. Molecular biology and evolution, 32, 268–274.

