## Supplementary Information for "Cryptic genetic variations of alanine:glyoxylate aminotransferase shape its fitness and dynamics"

**Table S1. Data collection and refinement statistics**

| **Data collection** | **AGT-Mi** |
|---|---|
| Beamline | Elettra XRD1 |
| Space group | $P4_1$ |
| Cell dimension; a b c (Å), α β γ (˚) | 90.6 90.6 140.9, 90 90 90 |
| Resolution range (Å) | 64.06 – 2.20 (2.93 - 2.20) |
| $R_{merge}$ | 0.092 (0.298) |
| Half-set correlation CC(½) (%) [(b)] | 99.2 (78.8) |
| <I/σI> | 9.3 (3.6) |
| Completeness (%) | 99.6 (99.5) |
| N. Reflection (total) | 214921 (16842) |
| Multiplicity | 3.8 (3.8) |
| Wilson B-factor (Å$^2$) | 23.8 |
| **Refinement** | |
| Resolution range (Å) | 64.0 - 2.20 |
| N° of unique reflections | 57240 |
| $R_{free}$ test set | 5660 refections (9.89%) |

| | |
|---|---|
| $R_{work}/R_{free}$ (%) | 23.9/28.9 |
| $F_o,F_c$ correlation | 0.88 |
| Average B-factor all atoms (Å$^2$) | 29.0 |
| Bonds RMSD | |
| *Length (Å)* | 0.074 |
| *Angle (°)* | 1.589 |
| Ramachandran plot (n. residues, %) | |
| *Favoured*<br>*Allowed* | 639 (96.0)<br>24 (4.0) |
| *(a) Values in parentheses refer to the highest-resolution shell*<br>*(b) Percentage of correlation between intensities from random half-data sets* | |

**Table S2. Molecular Dynamics parameters**

| parameter | value | comment |
|---|---|---|
| title | production MD simulation | |
| | | **Run parameters** |
| integrator | md | leap-frog integrator |
| nsteps | 100000000 | 2 * 100000000 = 20000000 ps (200 ns) |
| dt | 0.002 | 2 fs |
| | | **Output control** |
| nstenergy | 5000 | save energies every 10.0 ps |
| nstlog | 5000 | update log file every 10.0 ps |
| nstxout-compressed | 5000 | save coordinates every 10.0 ps |
| | | **Bond parameters** |
| continuation | yes | continuing from NPT |
| constraint_algorithm | lincs | holonomic constraints |
| constraints | h-bonds | bonds to H are constrained |
| lincs_iter | 1 | accuracy of LINCS |
| lincs_order | 4 | also related to accuracy |
| | | **Neighbor searching and vdW** |
| cutoff-scheme | Verlet | |

| | | |
|---|---|---|
| ns_type | grid | search neighboring grid cells |
| nstlist | 20 | largely irrelevant with Verlet |
| rlist | 1.2 | |
| vdwtype | cutoff | |
| vdw-modifier | force-switch | |
| rvdw-switch | 1 | |
| rvdw | 1.2 | short-range van der Waals cutoff (in nm) |
| | | **Electrostatics** |
| coulombtype | PME | Particle Mesh Ewald for long-range electrostatics |
| rcoulomb | 1.2 | |
| pme_order | 4 | cubic interpolation |
| fourierspacing | 0.16 | grid spacing for FFT |
| | | **Temperature coupling** |
| tcoupl | V-rescale | modified Berendsen thermostat |
| tc-grps | Protein non-Protein | two coupling groups - more accurate |
| tau_t | 0.1 0.1 | time constant, in ps |
| ref_t | 310 310 | reference temperature, one for each group, in K |
| | | **Pressure coupling** |
| pcoupl | Parrinello-Rahman | pressure coupling is on for NPT |
| pcoupltype | isotropic | uniform scaling of box vectors |
| tau_p | 2.0 | time constant, in ps |
| ref_p | 1.0 | reference pressure, in bar |
| compressibility | 4.5e-5 | isothermal compressibility of water, bar^-1 |
| | | **Periodic boundary conditions** |
| pbc | xyz | 3-D PBC |
| | | **Dispersion correction is not used for proteins with the C36 additive FF** |

| | | |
|---|---|---|
| DispCorr | no | |
| | | **Velocity generation** |
| gen_vel | no | continuing from NPT equilibration |

**Table S3 (provided as separate file). Multiple sequence alignment obtained using MUSCLE on Jalview program of the sequences retrieved from the non-redundant database using Blast.**

**Table S4. Sequence of primers used for the site-directed mutagenesis**

| **Mutation** | **Primer sequence (5'-3')** |
|---|---|
| L101A | GGACTCCTTCGCGGTTGGGGCCAATG |
| H146A | CCTGGCCCAGGCGAAGCCAGTGCTGCTG |
| L172A | TCGGGGAAGCGTGCCACAGGTACAAGTGCC |
| C173A | CGGGGAACTCGCGCACAGGTACAAGTGC |

**Table S5. Sequence of primers used for library generation**

| **Primer** | **Primer sequence** |
|---|---|
| AGT_vector_fw | CCTGCAGCACTGCCCCAAGAAGAAGCTGTG |
| AGT_vector_rv | GGGGGTCACCAGCAGCTTGTGAGAGGC |
| AGT_1_frag_fw | CAAGCTGCTGGTGACCCCCCCCAA |
| AGT_1_frag_rv | GCCCCCAAATGCCATTGGCCCCAAC |
| AGT_2_frag_fw | GGGCCAATGGCATTTGGGGGCAGCG |
| AGT_2_frag_rv | CATCAAGGGGCTGCAGCACGCCG |
| AGT_3_frag_fw | GCGTGCTGCAGCCCCTTGATGGCTTCG |
| AGT_3_frag_rv | GCAGTGCTGCAGGGCCGCCC |
| AGT_1_frag_fw | CAAGCTGCTGGTGACCCCCCCCAA |
| AGT_1_frag_rv | GCCCCCAAATGCCATTGGCCCCAAC |
| S99A-WT_RV_1 | CAACCAGGAAGGMGTCCCCAGGCTCC |
| E129K+G132E/G_fw_2 | GTGCACCCGATGACCAAGAAACCTGGAGRRCACTACAC ACTGC |

| | |
|---|---|
| R122al_P125al_T127al_fw_2 | GCATAGGAGCCCRSGTGCACCVGATGRYCAAGGACCCTGG |
| Pro125al-Thr127al_rv_2 | TCCAGGGTCCTTGRYCATCBGGTGCA |
| E138D/WT_RV_2 | CAGGCCCTCCTCCACMTCCTGCAGTGTGTA |
| A144T/WT+Q145R/WT_FW_2 | GGAGGGCCTGRCCCRGCACAAGCCAGTG |
| E171A/WT_FW_3 | CCCTTGATGGCTTCGGGGMACTCTGCCACAG |
| Y176H/WT_RV_3 | CCAGGAGCAGGCACTTGTRCCTGTGGCAGA |
| K177N/WT_FW_3 | CTCTGCCACAGGTACAASTGCCTGCTCCTGGTGGA |
| K177Q/WT_FW_3 | CTCTGCCACAGGTACMARTGCCTGCTCCTGGTGGA |
| E171A/wt-K177N/wt_FW_3 | GGGMACTCTGCCACAGGTACAASTGCCTGCTCCTGGTGGA |

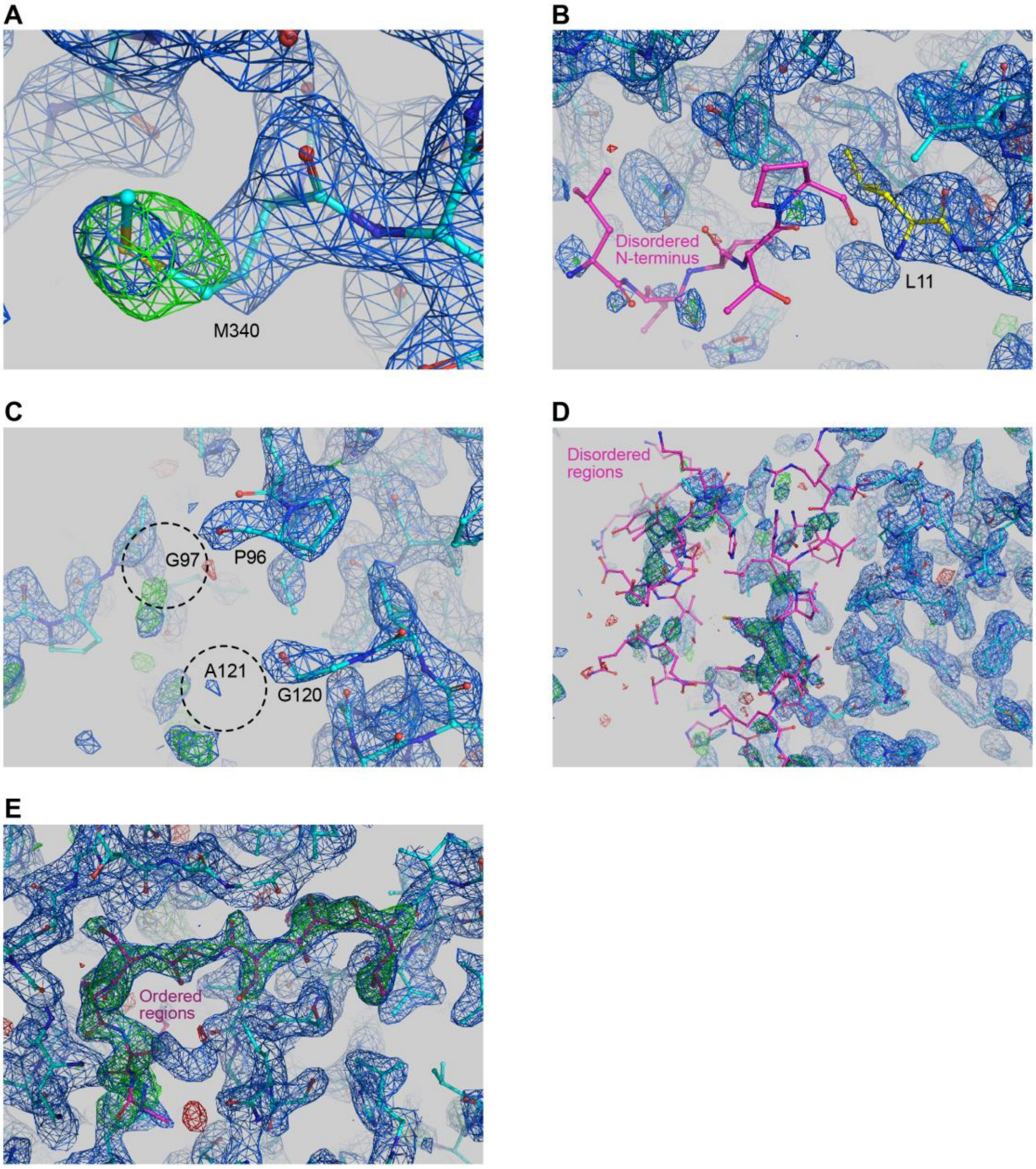


**Fig. S1 Electron density maps.** A) Electron density of maps after refinement of a model with residue 340 corresponding to an alanine. The final model (M340) is shown in cyan. 2Fo-Fc and Fo-Fc maps are contoured at 1.5 and 3.5 sigma, respectively. B) 2Fo-Fc and Fo-Fc maps (contoured as in panel A) of the N-terminus region. The first residue that could be modelled with confidence in the electron density was L11 (yellow), the position of the residues shown in magenta was not clear and therefore they were excluded from the final model. C and D) Maps (contoured as in panel A) of the disordered regions 1 and 2 (residues 97-102 and 121-146, respectively) are shown as an example of uninterpretable density due to multiple conformations. When the disordered regions were removed from the refined model, no density was present to build the missing residues, as shown for G97 and A121. E) Example of interpretable electron density maps, obtained by omitting residues 203-211 from the model. A clear electron density is present for the omitted section. The missing residues are shown in magenta.

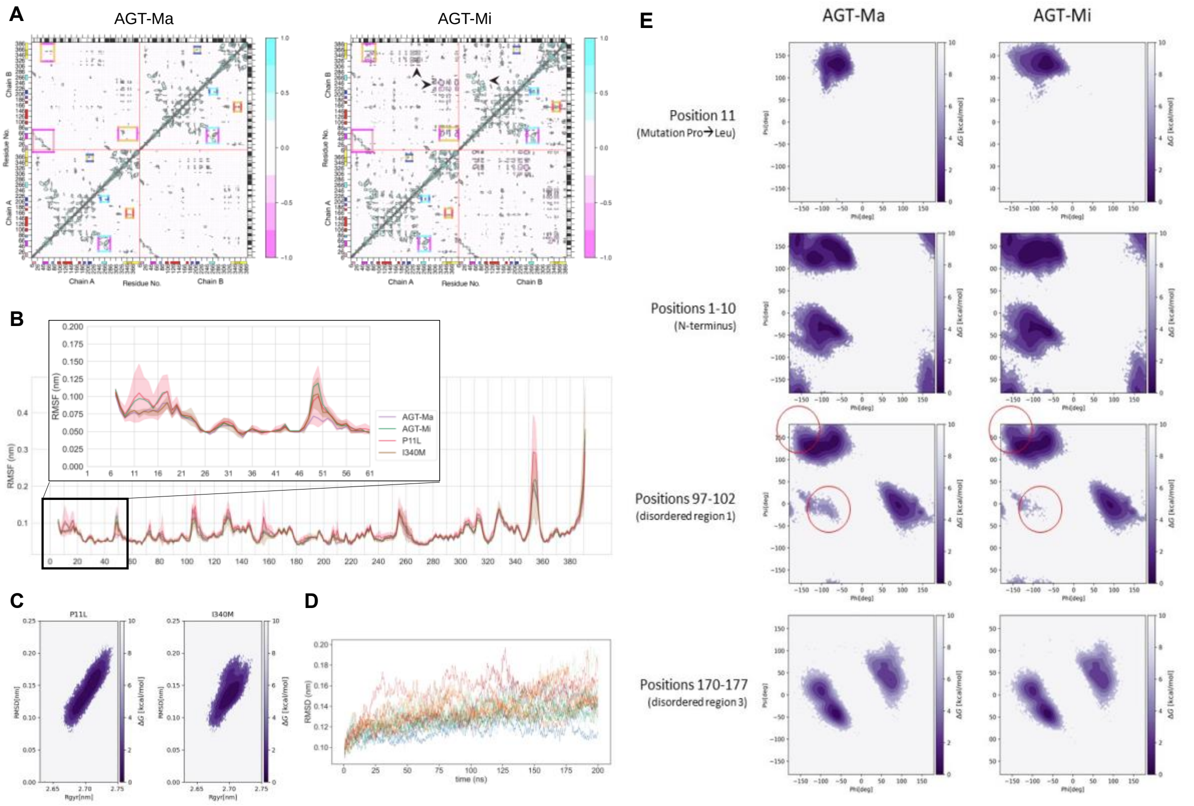


**Fig. S2 Molecular dynamics of AGT alleles and single mutants.** A) Dynamical cross-correlation matrix (DCCM) representing positive or negative correlation of motion obtained from the five joint repeats of AGT-Ma and AGT-Mi simulations. The color code of the annotation follows the one used in Fig 2A. The plot was obtained using the Bio3D R package (1) B) The chain and repeat averaged root mean square fluctuations (RMSF) of AGT single mutants P11L and I340M in comparison to the two main alleles. A Zoom-in of the first 60 positions is shown in the inlet. C) Free Energy Surface (FES) of the single mutant simulations. Root mean square deviation and the radius of gyration were used as coordinate systems. D) The root mean square deviation from the starting structure of each simulation is plotted against time. AGT-Mi, AGT-Ma, P11L and I340M are respectively represented in shades of blue, green, red and orange. The simulations quickly converge within 0.1 nm of deviation. E) Free energy landscape as a function of the backbone dihedral angles ($\Phi$ and $\Psi$) of several regions (7-10, 97-102, 170-177 and residue 11 alone). The angles were collected from five repetitions of 200 ns MD simulations of AGT-Ma and AGT-Mi structures. The same plot for disordered region 2 is shown in the main text (Fig.2B). Differences are observed for disordered region 1 (97-102), forming a β-sheet with region 2 and, as expected, for the mutated residue at position 11.

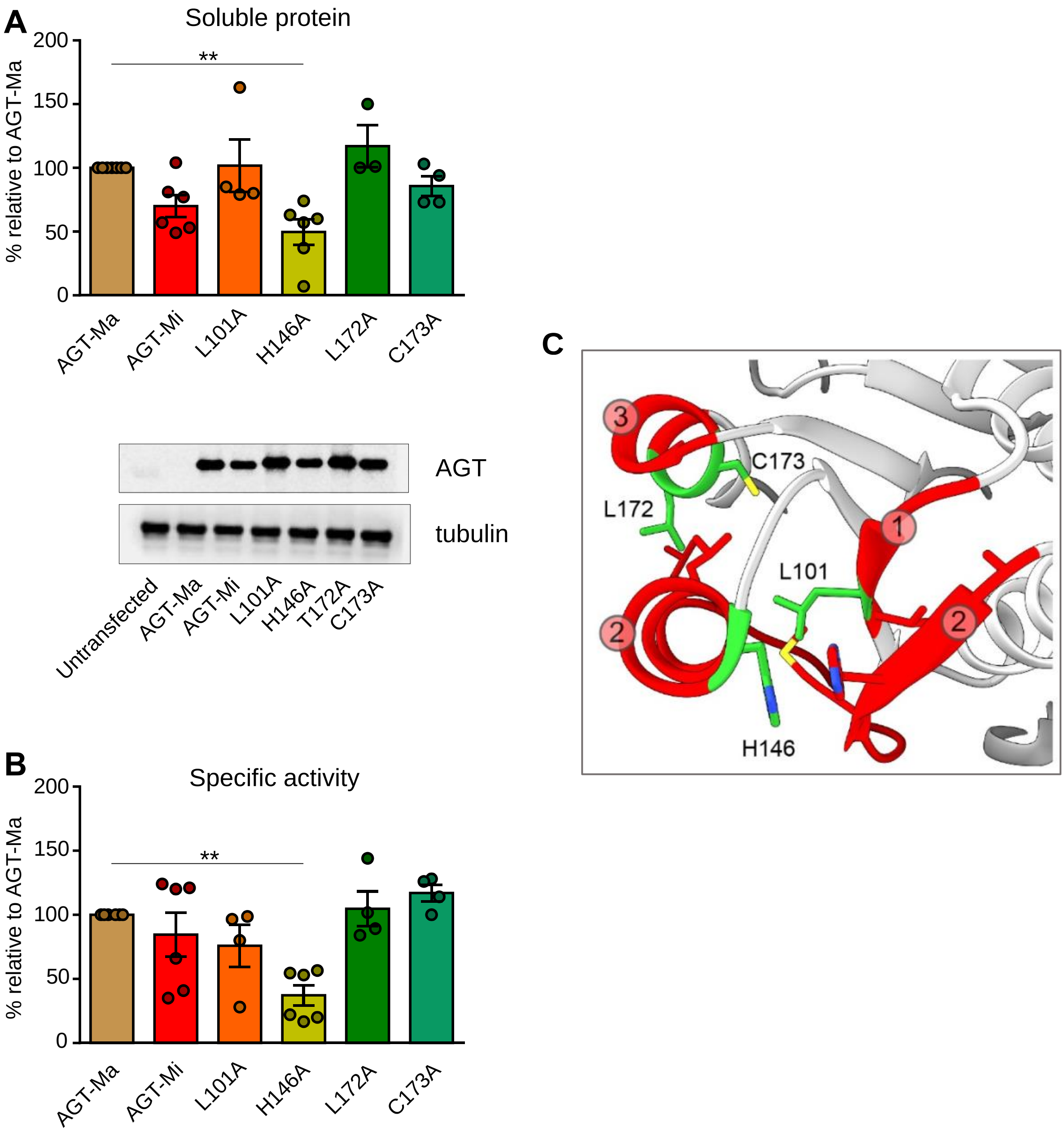


**Fig. S3 Expression and activity of AGT1 variants showing mutations of residues in regions prone to disorder.** HEK293 cells (panels A-B) were transiently transfected with pcDNA3.1 vectors encoding the indicated species. Cells were harvested and lysed. A) 10 μg of soluble lysate were subjected to SDS-PAGE, immunoblotted with anti-AGT from rabbit (1:10000), and detected by chemiluminescence. The histograms are representative of immunoblot band volume (as mean ± SEM of at least three independent experiments). B) Specific activity for the transamination reaction was determined by incubating 90 μg of soluble lysate with 0.5 M L-alanine, 10 mM glyoxylate, and 200 μM PLP at 25 °C in 100 mM KP, pH 7.4. C) Position of the mutated residues (green) in the three dynamic regions (residues 97-102=1; 120-146=2; 170-177=3). Mutants L110A and H146A destabilise the interaction between IDRs 1 and 2 (i.e. an hydrophobic interaction and an aromatic stacking interaction), while L172A and C173A destabilise the hydrophobic interactions between regions 2 and 3.

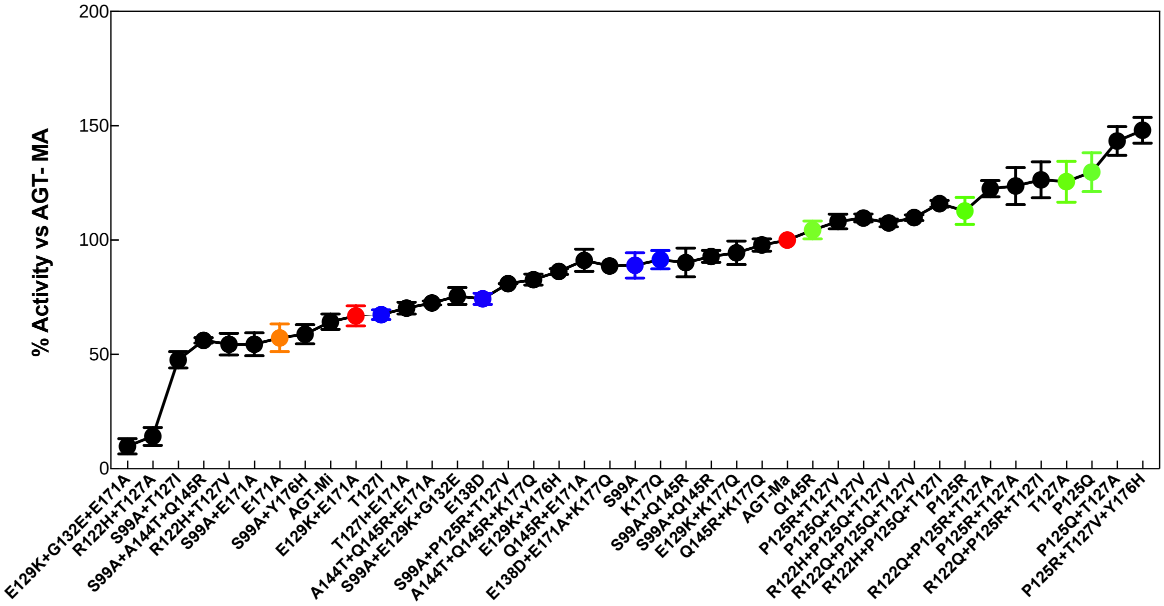


**Fig. S4. Specific activity of the variants obtained from the screening of the library.** All constructs have been expressed in *E.coli* cells and the specific activity for the transamination reaction has been measured using a spectrophotometric assay coupled with lactate dehydrogenase. AGT-Ma and AGT-Mi are represented as red dots while the single mutants are coloured based on their specific activity (orange, below AGT-Mi; blue, between AGT-Mi and AGT-Ma, and green above AGT-Ma). Data are represented as mean values ± SD. Experiments have been performed in duplicate and for each single experiments three technical triplicates have been used.

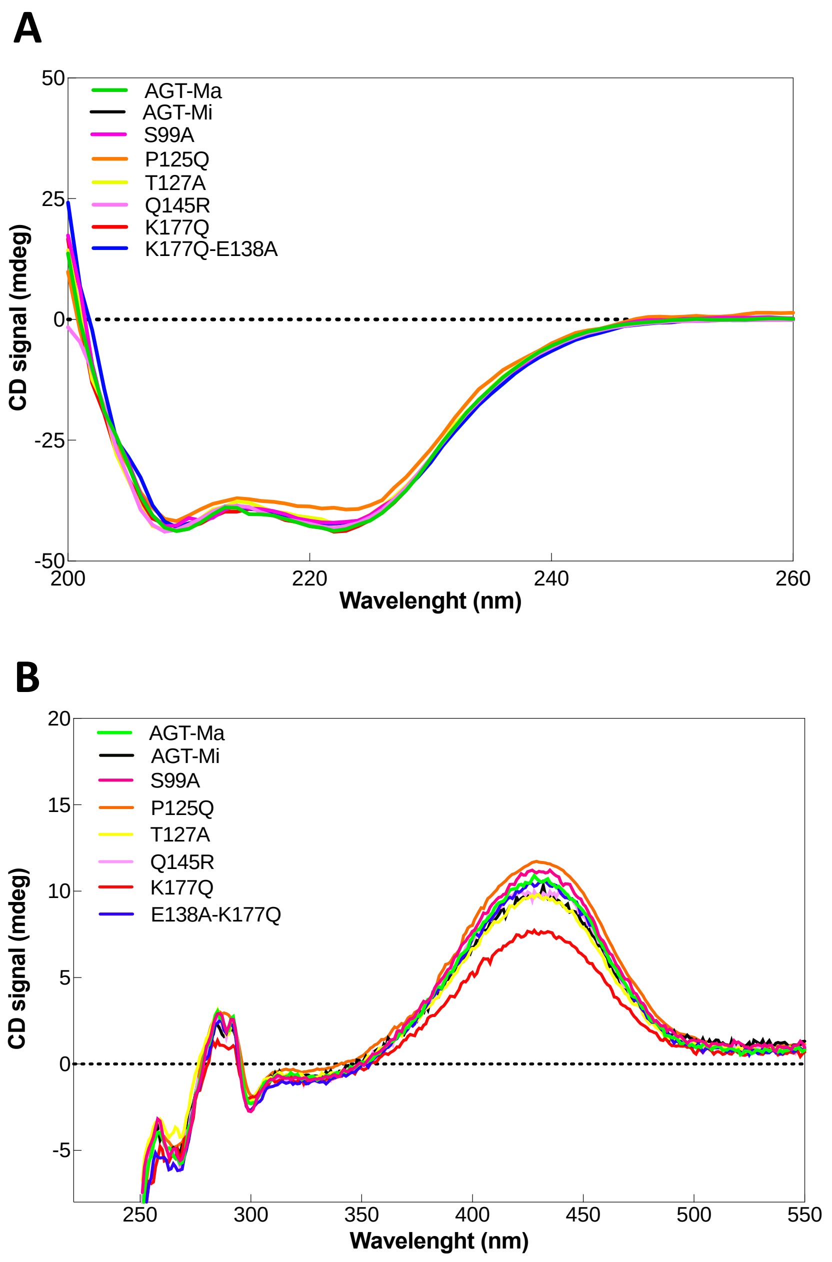


**Fig. S5. Far-UV (panel A) and Near-UV (panel B) CD spectra of AGT-Ma, AGT-Mi and the purified variants chosen from the library.** Protein concentration used was 8-10 μM and analyses have been performed in the presence of 20 μM exogenous PLP in KP 0.1 M pH 7.4 at 25 °C. All the spectra have been recorded using a Jasco J-820 spectropolarimeter (see details in the Materials and Methods).

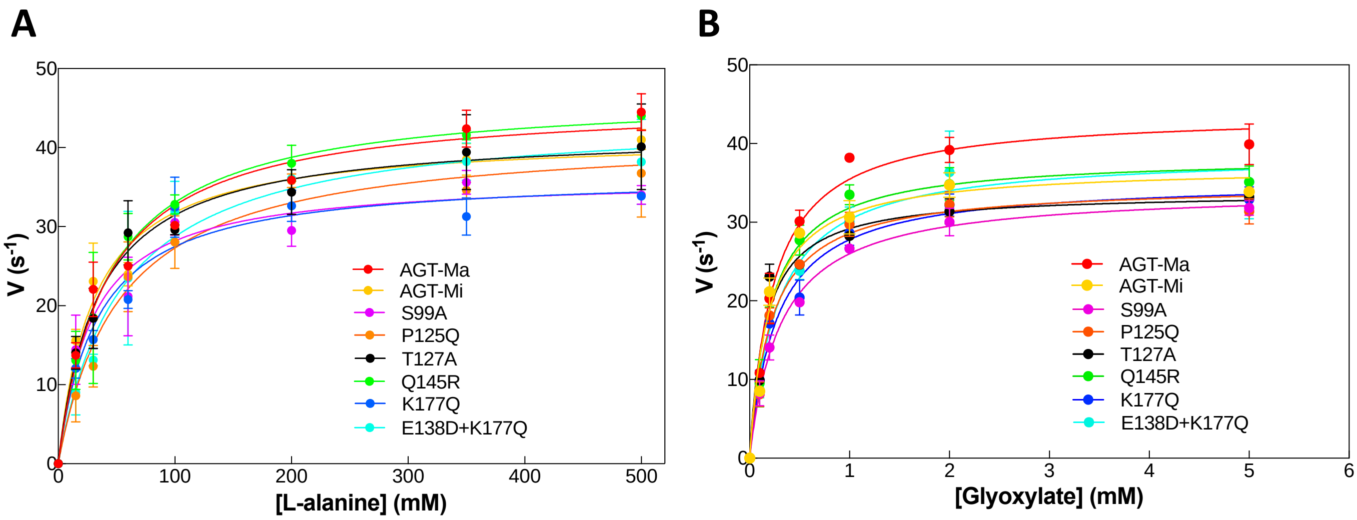


**Fig. S6. Michaelis-Menten curves of AGT-Ma, AGT-Mi and the variants selected from the library.** A) Values of initial velocity (nmol product/time/nmol enzyme) as a function of L-alanine concentration (15-500 mM) in the presence of 10 mM glyoxylate. B) Values of initial velocity (nmol product/time/ nmol enzyme) as a function of glyoxylate concentration (0.07-5 mM) in the presence of 500 mM L-alanine. The enzyme concentration was 100-200 nM and assays were performed in the presence of 50 μM exogenous PLP in KP 0.1 M pH 7.4 at 25 °C. Data are represented as mean values ± SD of three independent experiments (n = 3).

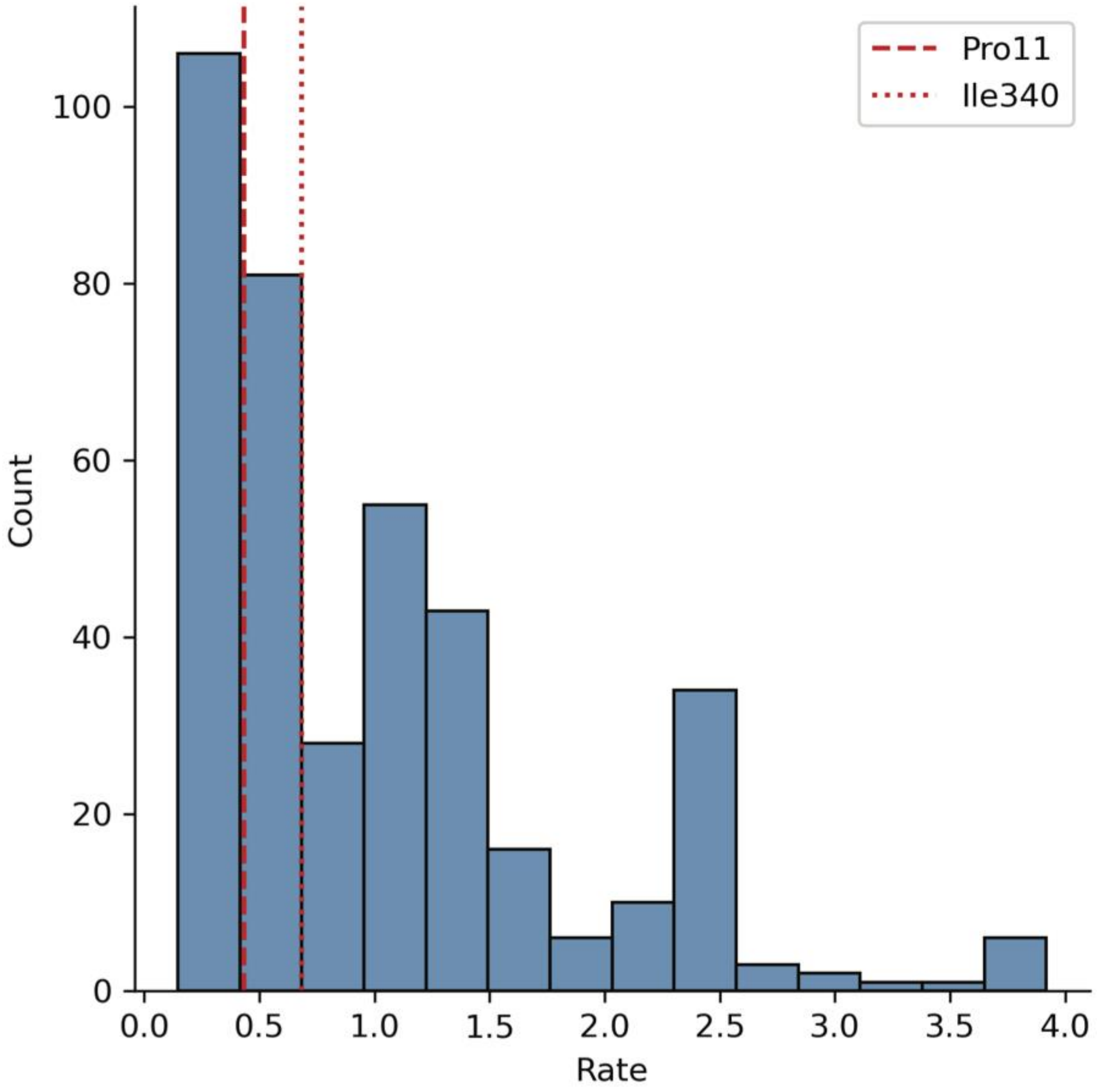


**Fig. S7. Distribution of the per-site rate of evolution of AGXT1 vertebrate phylogeny.** The rate of evolution per-site was calculated with IQTREE and plotted as a distribution. The values represent the posterior mean site rate weighted by posterior probability (2). The two red lines indicate the rate of evolution of the sites of AGT-Ma that are mutated in the AGT-Mi.

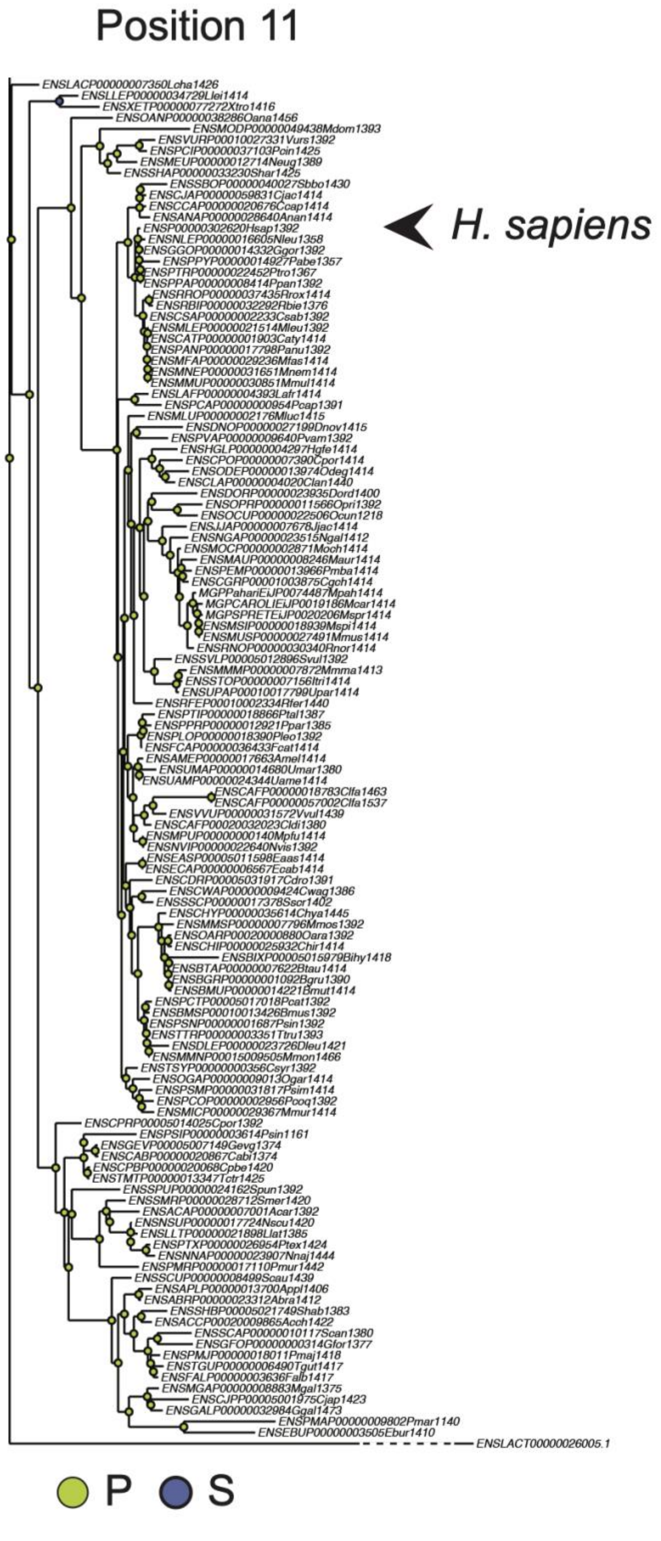


**Fig. S8. Phylogenetic tree of AGXT1 protein sequences annotated by position 11 ancestor.** The phylogenetic analysis of 254 AGXT1 vertebrate genes was annotated with the ancestor sequence of position 11 using PastView (3). The fish clade was cut from the picture, because all the nodes show P as ancestral amino acid.

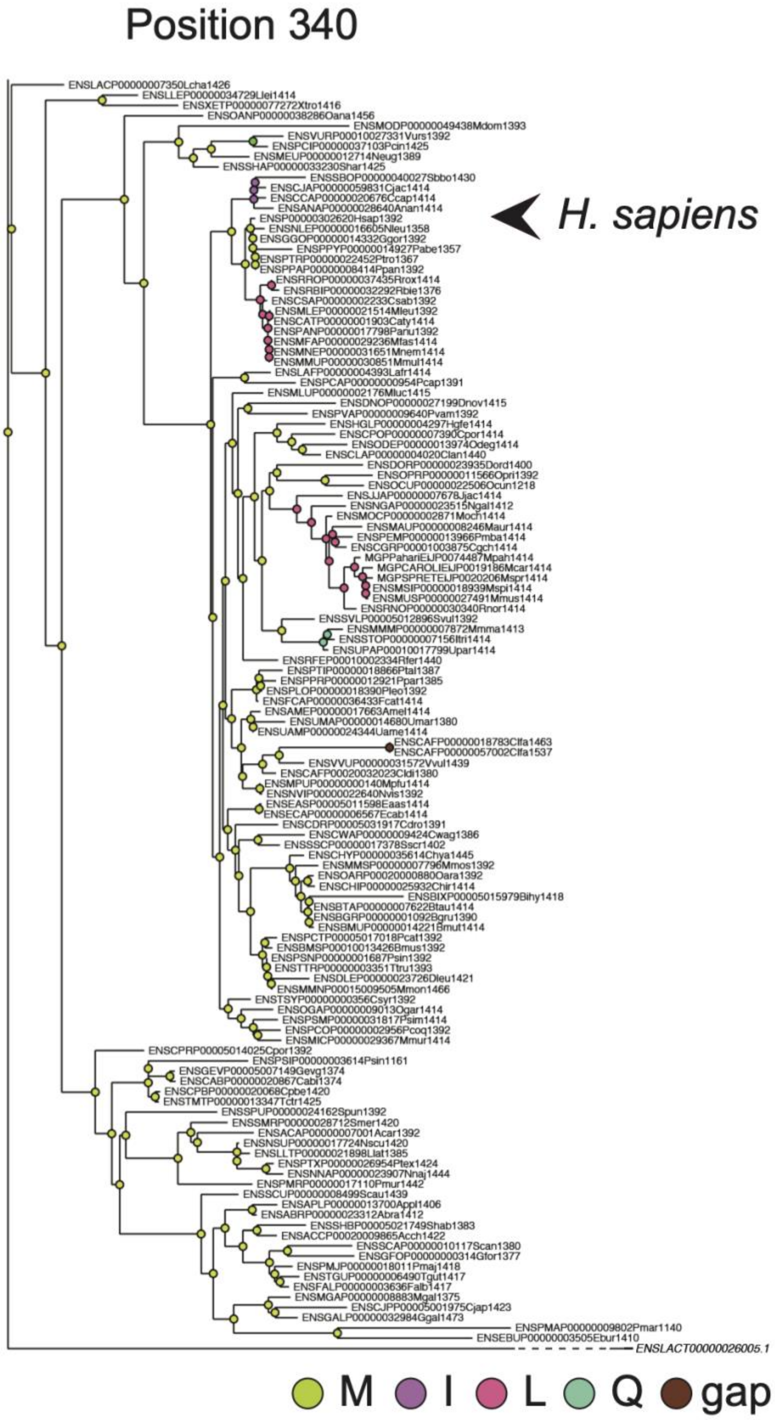


**Fig. S9. Phylogenetic tree of AGXT1 protein sequences annotated by position 340 ancestor.** The phylogenetic analysis of 254 AGXT1 vertebrate genes was annotated with the ancestor sequence of position 340 using PastView (3). The fish clade was cut from the picture, because all the nodes show M as ancestral amino acid.

**Legend to Movie S1: AGT-Ma conformational frustration.** A complete rotation around the dimer's 2-fold axis shows how frustrated residues connect the regions that fluctuate more. For a complete description of this structural representation see the legend of Figure 2B.

### REFERENCES


1 Grant, B.J., Rodrigues, A.P.C., ElSawy, K.M., McCammon, J.A. and Caves, L.S.D. (2006) Bio3d: an R package for the comparative analysis of protein structures. *Bioinformatics*, **22**, 2695-2696.

2 Mayrose, I., Graur, D., Ben-Tal, N. and Pupko, T. (2004) Comparison of site-specific rate-inference methods for protein sequences: empirical Bayesian methods are superior. *Molecular biology and evolution*, **21**, 1781-1791.

3 Chevenet, F., Castel, G., Jousselin, E. and Gascuel, O. (2019) PastView: a user-friendly interface to explore ancestral scenarios. *BMC Evolutionary Biology*, **19**, 163.
